# Compressive mechanical stress activates ERK5 to regulate cortical tension and promote invasive cellular traits

**DOI:** 10.64898/2026.01.30.702739

**Authors:** Nevena Srejic, Frank van Drogen, Tatjana Sajic, Federico Ulliana, Matthias Peter

## Abstract

Growth-induced compressive forces regulate diverse biological processes ranging from development to tumor progression, but the underlying molecular mechanisms remain poorly understood. Here, we found that compressive mechanical cues activate the MAP kinase extracellular signal-regulated kinase 5 (ERK5) at the cell periphery, inducing invasive traits in epithelial cells. Using a phospho-proteomic approach combined with an RNAi-based validation screen, we identified several previously unrecognized ERK5 effectors important for cell-cell adhesion and actin organization, among them the myosin light chain (MLC) phosphatase subunit Myosin Phosphatase-Targeting Subunit 1 (MYPT1). Our results demonstrate that ERK5 regulates MYPT1, which in turn inhibits myosin at the cell periphery. ERK5 signaling counteracts Rho-associated protein kinase (ROCK), thereby reducing peripheral actomyosin tension. This relaxation destabilizes cell-cell contacts and promotes cell migration in response to mechanical compression. Together, these findings identify an ERK5-mediated response mechanism activated by compressive force that triggers cell invasion, highlighting ERK5 therapeutic potential.

## Introduction

Cells convert mechanical forces into biochemical signals through the processes of mechanosensation and mechanotransduction, dynamically adapting their behavior and function to their surroundings^1^. For instance, cells must sense and respond to shear stress, tension, compression, and extracellular matrix (ECM) stiffness to maintain tissue homeostasis^2^. Cells experience compressive mechanical forces when they are physically pressed against neighboring cells or surrounding tissue components, owing to increases in cell and matrix density^3^. These forces are inherent to normal physiology such as orchestrating tissue patterning during embryogenesis and organogenesis^4,5^. Compressive forces are also linked to disease states^6–8^. For example, the expanding tumor mass generates forces directed from the tumor core toward the surrounding tissue, while the remodeled host tissue generates inwardly projecting forces resisting tumor expansion^9–11^. As a result, both cancer cells and adjacent normal cells are subjected to altered physical inputs that can influence their behavior and function^10^. Despite the patho-physiological significance and strong correlation with cancer aggressiveness^3,6^ and therapeutic resistance^3^, the molecular mechanisms and signaling networks governing mechano-transduction of compression in mammalian systems remain largely unknown.

Prior studies in yeast found that compressive forces activate the Cell Wall Integrity (CWI) pathway^12^, with its component Mitogen-Activated Protein Kinase 1 (Mpk1) being an ortholog of extracellular signal-regulated kinase 5 (ERK5)^13^. ERK5 has structural distinctions from other canonical MAP kinases, with only a few physiological substrates identified to date^14^. Interestingly, laminar shear stress (LSS) was shown to activate ERK5, conferring a cytoprotective effect on vascular endothelial cells^15,16^, while in mammary gland epithelial cells ERK5 was found to be involved in TGFβ-induced epithelial-mesenchymal transition (EMT)^17^. During EMT epithelial cells lose their cell-to-cell adhesion and polarity, resulting in transformed cells that can migrate and invade other tissues^17,18^. While crucial for development, EMT can also be involved in diseases like fibrosis and cancer metastasis^18^. Indeed, aberrant expression and activation of ERK5 and its upstream regulator, Mitogen-Activated Protein Kinase Kinase 5 (MEK5), have been reported in several cancer types, including breast cancer, correlating with the acquisition of more aggressive tumor phenotypes^19–22^.

Here, we identify the MAP kinase ERK5 as a critical signaling mediator of compressive stress response. Indeed, compressive stress leads to rapid ERK5 activation at the periphery of Normal Murine Mammary Gland Epithelial (NMuMG) cells, and ERK5 is required for invasive cellular features, including rearranging the cortical actin cytoskeleton and disassembling cell-cell junctions. Our results demonstrate that upon compression, ERK5 regulates the myosin light chain (MLC) phosphatase subunit MYPT1 to inhibit myosin contraction, thereby dissociating cell-cell contacts and promoting cell migration.

## Results

### Compressive mechanical stress triggers morphological rearrangements in NMuMG cells

The mammary gland is a highly mechanoreactive tissue that can undergo repetitive rounds of distinct physiological changes, such as epithelial-mesenchymal transition during development, homeostasis, and pathological conditions^23,24^. For example compressive forces become particularly important during milk stasis and breast cancer progression when luminal epithelial cells are pushed against myoepithelium and basement membrane either due to milk accumulation or uncontrolled cell proliferation^11,25^. We thus used Normal Murine Mammary Gland Epithelial (NMuMG) cells as a physiological model to study compressive stress signaling. We adapted a transwell-based platform to apply compressive stress mimicking forces at tissue interfaces (Fig.1a)^6^. Cells were grown on a transwell insert with 0.4 µm or 8.0 µm pores in serum-rich medium, and then serum-starved for 24 h to minimize effects of cell proliferation and growth factor signaling (Extended Data Fig. 1a). An agarose cushion was placed on top of the cells and compressive stress was applied via custom-built metal weights. The load exerts a uniaxial pressing force of 4 mmHg, which falls within the range of compressive stress measured in solid tumors^26,6^.

**Fig. 1.**
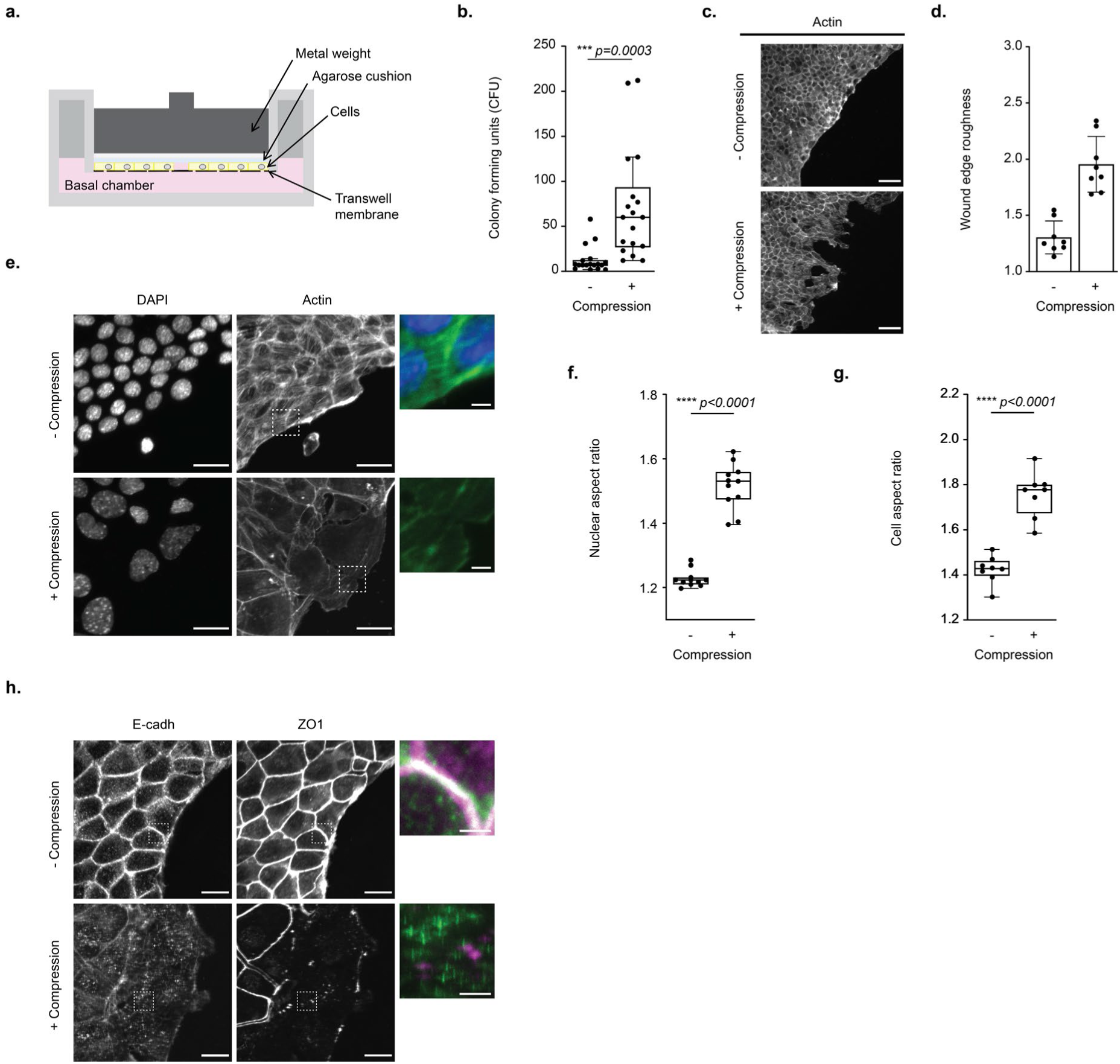
Compressive mechanical stress triggers an invasive response and morphological rearrangements of NMuMG cells. **a**. Schematic drawing of the transwell-platform. Serum deprived NMuMG cells were covered with an agarose disk followed by a metal load of defined weight, exerting uniaxial compressive force. At the indicated times, the load was removed, and the cells were harvested for different assays. **b**. NMuMG cells were exposed (+) or not (-) to compression and allowed to migrate for 24 hours through 8 µm transwell pores. Transmigrated cells adhering to the bottom of the 6-well plate form colonies, which were stained with crystal violet. The colony-forming units were quantified to compare the migratory potential across conditions. N=18; unpaired t test comparison; p<0.0001****; p<0.05 considered significant. **c**. and **d**. Serum deprived NMuMG cells invading the empty space were imaged by phalloidin staining 24 hours after removing the inset of the ibidi chambers, in the presence (+) or absence (-) of compressive pressure. Scale bar: 50 μm (**c**). Note that compressed cells acquire an elongated, migratory-like phenotype, due to a profound rearrangement of their cytoskeleton. The roughness of the edge was quantified (**d**) as described in Extended Data Fig. 1c. The normalized wound edge perimeter was plotted as a metric of the compressive stress phenotype. Unpaired t-test comparison; p<0.0001****; p<0.05 considered significant. **e**, **f** and **g**. NMuMG cells under normal (-) or compressive stress (+) conditions were stained with DAPI (grey and blue) or rhodamine-phalloidin (grey and green), to visualize nuclei or the actin cytoskeleton, respectively (**e**). Scale bar: 20 μm. The inset magnifies a section of cells at the epithelia edge. Scale bar: 3 μm. The nuclear (**f**) and cell (**g**) shape changes were quantified as an increase in the cell or nuclear aspect ratio after cellpose-based cell or nuclear segmentation. N=11 and N=8, respectively; Unpaired t-test comparison; p<0.0001****; p<0.05, considered significant. **h**. The adherens junction marker E-cadherin (gray and green) and the tight junction protein ZO1 (gray and magenta) were immunostained in compressed (+) and non-compressed (-) NMuMG cells. Scale bar: 20 μm. The inset magnifies a section of cells at the epithelia edge. Note that upon compression, E-cadherin is internalized and forms punctate structures, while ZO1 disappears from tight junctions.

Interestingly, NMuMG cells compressed for 24 h readily migrated through 8.0 μm pores, as measured by the increasing number of colony-forming units (CFU) in the basal chamber (Fig. 1b, Extended Data Fig. 1b). To investigate this invasive phenotype, we seeded NMuMG cells on 0.4 µm membranes in Ibidi inserts, creating two regions with cells separated by a cell-free gap, which allows observing compressed cells when they invade the wound. We first visualized nuclear morphology, since nuclear deformation is an integral part of mechano-responses^27,28^. As expected, upon compression, cell nuclei exhibited an elongated morphology reflected by an increased nuclear aspect ratio (Fig. 1e, f; Extended Data 1d), confirming that cells experience and respond to the applied mechanical force. Staining actin structures by rhodamine-phalloidin revealed that compressed NMuMG cells acquire an invasive, irregular tissue front, while the edge of uncompressed cells remained smooth (Fig. 1c). We quantified the edge roughness as the perimeter of the tissue edge divided by the hypothetical perfectly smooth edge between the two endpoints of the tissue line (Extended Data Fig. 1c). The edge roughness index significantly increased in compressed cells, which was caused by morphological changes in cells at the tissue edge (Fig. 1d) and not by increased cell proliferation of the quiescent cells. Indeed, compression leads to profound changes in the actin cytoskeleton of cells at the tissue edges, which form protrusions and disrupts junctional actin filaments (Fig. 1e). This striking elongation of edge cells was quantified and expressed as the aspect ratio (Extended Data Fig. 1d), which significantly increased upon compression (Fig. 1g). Since disruption of peripheral actin indicates dissociation of cell-cell contacts, we assessed cell-cell junctional integrity by staining E-cadherin and zonula occludens-1 (ZO1), to visualize adherens and tight junctions, respectively. Indeed, NMuMG cells lose both types of connections upon compression (Fig. 1h). While E-cadherin was internalized and observed in punctate cytoplasmic structures, ZO1 was dispersed in the cytoplasm of compressed edge cells. Taken together, we conclude that NMuMG cells respond to compressive cues by acquiring invasive properties at the tissue front characterized by distinct morphological rearrangements of the actin cytoskeleton and cell-cell junctions.

### Compressive stress activates ERK5 at the cell periphery

Since the MAP-kinase ERK5 was implicated in the cellular response to shear stress^15,16^ and is required for EMT^17^, we hypothesized that ERK5 might be an important signaling mediator of compressive stress in mammals. We thus measured transcriptional activity of myocyte enhancer factor 2 (MEF2) family members, the best-characterized ERK5 substrates^14^. We generated a transcriptional reporter construct containing MEF2-binding sites upstream of GFP, allowing analysis of ERK5-dependent GFP induction by western blot (Fig. 2a). Indeed, GFP expression was significantly increased in cells reverse-transfected with the reporter construct and exposed to compressive stress for 24 h (Fig. 2b). GFP expression was also increased in unstressed NMuMG cells co-transfected with constructs expressing HA-tagged constitutively-active MEK5 (HA-MEK5DD), which activates ERK5 by phosphorylating the conserved Thr-Glu-Tyr (TEY) motif (Fig. 2a). As expected, GFP induction was not observed in cells treated with the ERK5 kinase inhibitor XMD8-92 (Fig. 2b). Western blot analysis with a phospho-ERK5 specific antibody substantiated that compressive stress and HA-MEK5DD overexpression activate ERK5 (Fig. 2c, Extended Data 2a). Depletion of ERK5 by siRNA confirmed specificity of the ERK5 and phospho-ERK5 antibodies (Fig. 2c). We conclude that compressive mechanical stress activates ERK5, thereby enhancing MEF2-induced transcription. We next used immunofluorescence microscopy to visualize subcellular ERK5 activity in cells exposed to compressive stress. Strikingly, phospho-ERK5 accumulated at the cell periphery when compression was applied for 3 h (Fig. 2d), while no phospho-ERK5 signal could be detected in ERK5-depleted control cells. Using a cell segmentation pipeline, we quantified total fluorescence and peripheral intensity by defining a zone around the cell edge (Extended Data 2b). Indeed, both total and peripheral fluorescence intensity of phospho-ERK5 was increased upon compressive stress (Fig. 2e, f). Taken together, these results show that compressive cues activate peripheral ERK5 and suggest a function in cell-cell contact dissolution and actin rearrangements.

**Fig. 2.**
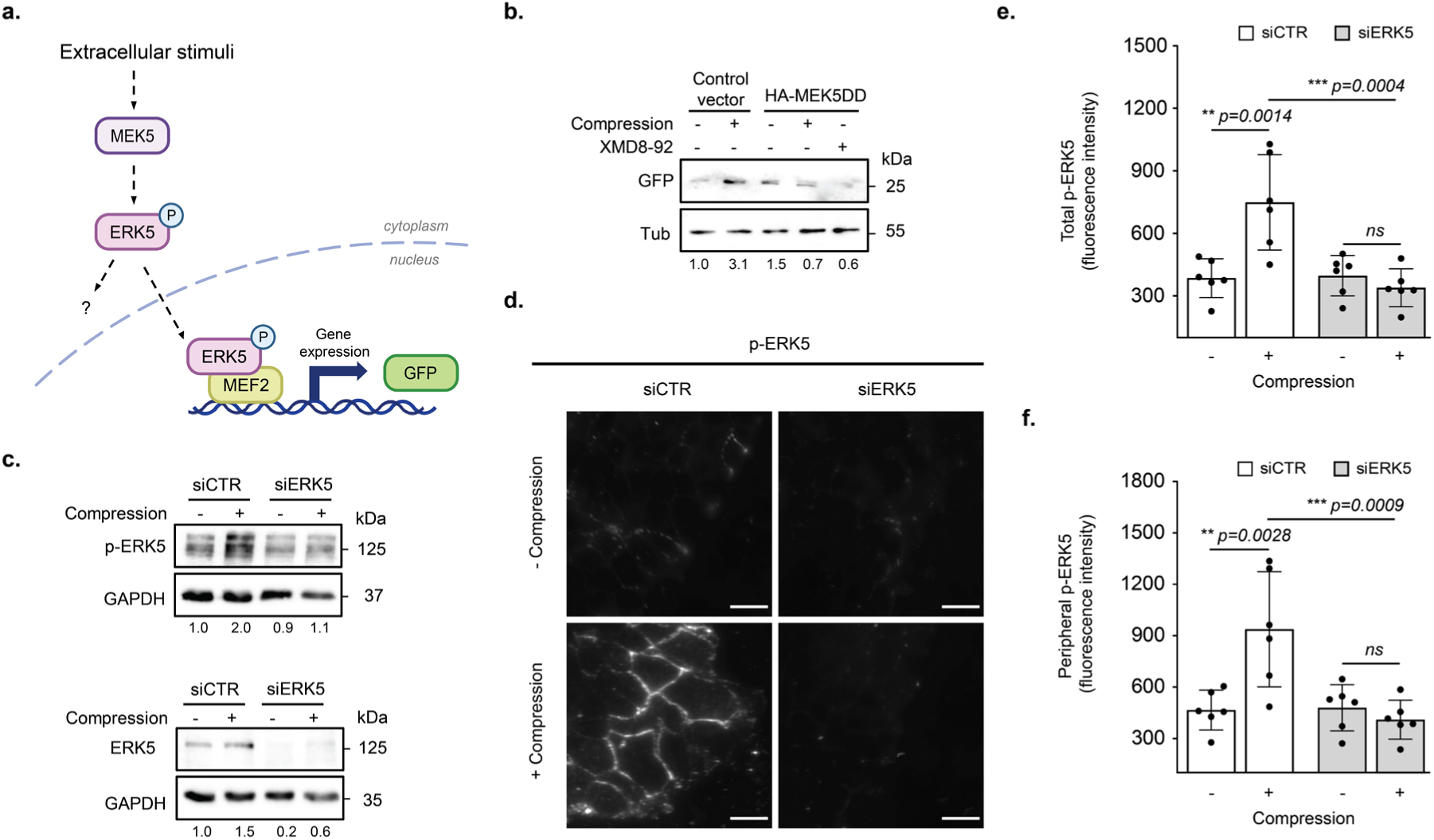
Compression leads to ERK5 activation and accumulation at cell-cell contact sites. **a.** Schematic representation of the ERK5 MAPK pathway. ERK5 is phosphorylated and thereby activated by MEK5 on the TY residues in its T-loop. Active ERK5 phosphorylates cytoplasmic targets, and translocates into the nucleus, where it binds and phosphorylates the transcription factor MEF2. In the reporter construct, active MEF2 drives expression of GFP (*MEF2p*-GFP). **b.** NMuMG cells were reverse-transfected with the *MEF2p*-GFP reporter plasmid and either an empty control vector (EV) or a vector expressing an HA-tagged dominant-active MEK5 mutant (HA-MEK5DD). GFP expression was monitored with (+) or without (-) compressive stress after 24 hours by western blot analysis. Where indicated (+), XMD8-92 was added to inhibit ERK5 activity. α-tubulin is used as a loading control. The numbers below represent densitometric quantification of GFP level across conditions, normalized to the tubulin loading control and relative to the empty control vector in the absence of compression and ERK5 inhibition. **c.** Western blot (WB) analysis confirmed elevated levels of active ERK5 (p-ERK5) after 3 hours of compression in serum-starved NMuMG cells, while the total ERK5 levels remained unchanged. α-tubulin controls equal loading. The numbers below represent the densitometric quantification of p-ERK5 and ERK5 levels across conditions normalized to GAPDH levels and relative to control condition without compression and ERK5 depletion. **d - f.** NMuMG cells siRNA-depleted for ERK5 (siERK5) or treated with a control siRNA oligo (siCTR) were exposed (+) or not (-) to compressive stress for 3 hours and the localization of p-ERK5 was analyzed by immunofluorescence analysis. Representative images show increased p-ERK5 at cell-cell contacts (**d**). Scale bar: 20 μm. Total- (**e**) and epithelial (**f**) p-ERK5 fluorescence intensity was quantified, and statistical significance confirmed by an ANOVA followed by Tukey multiple comparison test; ***p<0.001; p<0.05 was considered statistically significant.

### ERK5 is required for the invasive phenotype triggered by compressive cues

We next probed the functional importance of ERK5 when cells respond to compressive mechanical stress. Indeed, ERK5 siRNA depletion or inhibition of its catalytic activity with XMD8-92 significantly reduced colony-forming units in the transmigration assay (Fig. 3a, Extended Data 3a-c). Similarly, blocking ERK5 activity prevented increased edge roughness compared to controls (Fig. 3b-d, Extended Data 3d). Most notably, cells overexpressing HA-MEK5DD exhibited an invasive tissue edge, while the edge remained smooth in XMD8-92-treated cells or vehicle controls (Fig. 3e, f). Together, these data demonstrate that compression-mediated ERK5 activation is necessary and sufficient to induce an invasive phenotype in NMuMG cells. To corroborate these data, we used ERK5 siRNA depletion or XMD8-92 treatment to analyze the role of ERK5 in compression-mediated cellular responses, such as nuclear deformation, actin rearrangements, and resolution of cell-cell junctions. Interestingly, ERK5 activity in compressed NMuMG cells was required for edge cell elongation (Fig. 3h, Extended Data Fig. 3f) and actin cytoskeleton rearrangements (Fig. 3g, Extended Data Fig. 3e), including dissociation of peripheral actin bundles. Moreover, ZO1 maintained its localization when compression was applied in the absence of ERK5 activity (Fig. 3g, Extended Data Fig. 4a). Finally, E-cadherin internalization was significantly reduced and its peripheral signal maintained in the absence of ERK5 activity (Extended Data Fig. 4b, c). In contrast, nuclear elongation was not significantly affected by siERK5 or XMD8-92 treatment, and the nuclear aspect ratio stayed within the range of cells transfected with non-targeting siRNAs or treated with DMSO, respectively (Fig. 3g, i, Extended Data Fig. 3g). Conversely, HA-MEK5DD overexpressing NMuMG cells acquire elongated morphology characterized by actin protrusions and ruptured junctions. This phenotype was prevented when cells were treated with XMD8-92 (Extended Data Fig. 4d), suggesting that ERK5 activity is not only necessary but also sufficient to induce profound changes at the cell periphery. Taken together, these observations demonstrate that in response to compressive stress ERK5 orchestrates actin cytoskeleton reorganization and cell-cell contact disassembly, while nuclear changes require ERK5-independent mechanisms.

**Fig. 3.**
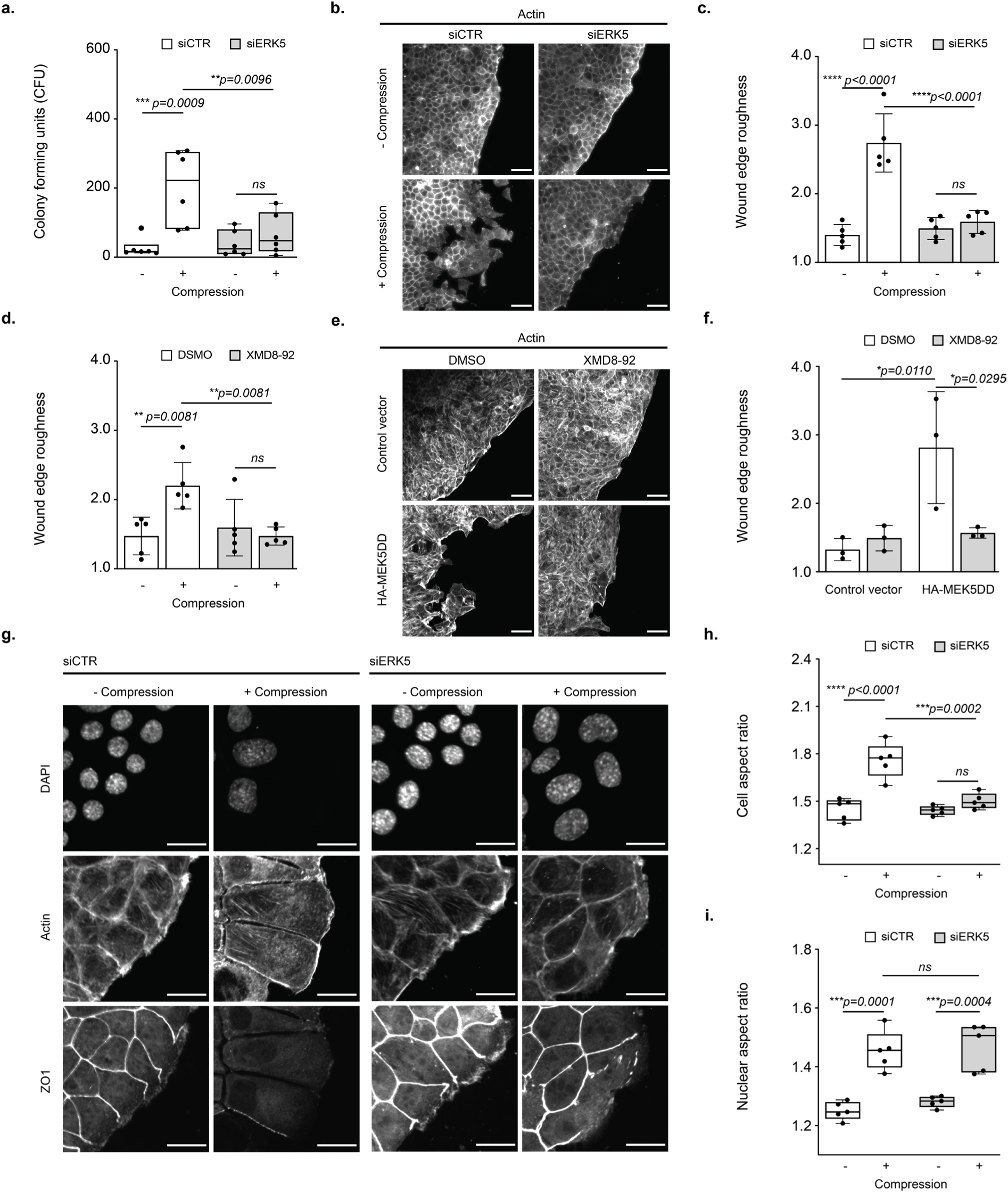
ERK5 activity is required to trigger the invasive phenotype of compressed NMuMG cells. **a.** NMuMG cells siRNA depleted for ERK5 (siERK5) or treated with a control oligo (siCTR) were exposed (+) or not (-) to physical compression and allowed to migrate for 24 hours through 8 μm transwell pores. Colonies formed by transmigrated cells were visualized by crystal violet staining and quantified as described in Figure 1b. N=6; ANOVA followed by Tukey multiple comparison test; ***p<0.001; p<0.05, considered statistically significant. **b - d.** Invading NMuMG cells growing in ibidi chambers, either siRNA-depleted for ERK5 or treated with control oligos (**b, c**) or exposed to the ERK5 inhibitor XMD8-92 or DMSO solvent control (**d**), were visualized by phalloidin staining in the presence (+) or absence (-) of compressive pressure for 24 hours. The wound edge roughness was quantified to compare the different conditions. N=5; ANOVA with Tukey multiple comparison; ***p<0.001; p<0.05, considered statistically significant. **e.** and **f.** NMuMG cells grown in ibidi chambers were reverse-transfected with an empty control vector or a vector expressing dominant-active HA-MEK5DD. Where indicated, the ERK5 inhibitor XMD8-92 or solvent control (DMSO) was added. The wound edge roughness was visualized by phalloidin staining (**e**) and quantified to compare the different conditions (**f**). N=3; ANOVA with Tukey multiple comparison; ***p<0.001; p<0.05, considered statistically significant. **g - i.** NMuMG cells siRNA-depleted for ERK5 (siERK5) or treated with control oligos (siCTR) exposed (+) or not (-) to compressive force conditions were stained with DAPI, rhodamine-phalloidin, and immunostained with antibodies against ZO1 to visualize tight junctions. Scale bar: 20 μm. The cell shape changes (**h**) and nuclear aspect ratio (**i**) were quantified as described in the legend to Figure 1f and g, respectively. N=5; ANOVA with Tukey multiple comparison; ***p<0.001; p<0.05, considered statistically significant.

### Activation of ERK5 triggers phosphorylation of several proteins that regulate the actin cytoskeleton and cell junctions

To identify relevant ERK5 substrates, we used mass-spectrometry for global proteome and untargeted phosphoproteomic analysis across different experimental conditions (Fig 4a). Few changes in protein abundance were detected (Extended Data Fig. 5a), demonstrating that the early compressive stress response is not primarily regulated by changes in protein expression or degradation. In contrast, clustering of the phosphoproteome data clearly separated the conditions exposed to pressure (Extended Data Fig. 5b and 5c) and allowed to distinguish sites phosphorylated by ERK5-dependent and ERK5-independent mechanisms (Fig. 4b-c). Interestingly, GO term analysis of compression-induced but ERK5-independent phosphorylation sites revealed significant enrichment of positive regulators of mRNA-mediated gene silencing, positive regulators of post-transcriptional gene silencing by RNA and RNA export from the nucleus (Fig. 4d). For example, this cluster contained RNA binding proteins regulating mRNA decay, splicing and trafficking (Fig. 4b), indicating that compressive stress triggers an ERK5-independent response of nuclear transport regulation and post-transcriptional gene silencing. Importantly, however, enrichment analysis revealed that compression-induced, ERK5-dependent biological processes involved proteins linked to cell-cell junctions, epithelial cell development and actin filament organization (Fig. 4e). For example, compression-induced and ERK5-dependent phosphorylation sites were found in proteins such as Filamin A, ZO1, ZO2, Zyxin, Myosin Phosphatase Target Subunit 1 (MYPT1) and LIM domain and actin binding 1 (Lima1), known structural components and regulators of the actin cytoskeleton and cell-cell contacts (Fig. 4c), consistent with the accumulation of active ERK5 at the cell periphery.

**Fig. 4.**
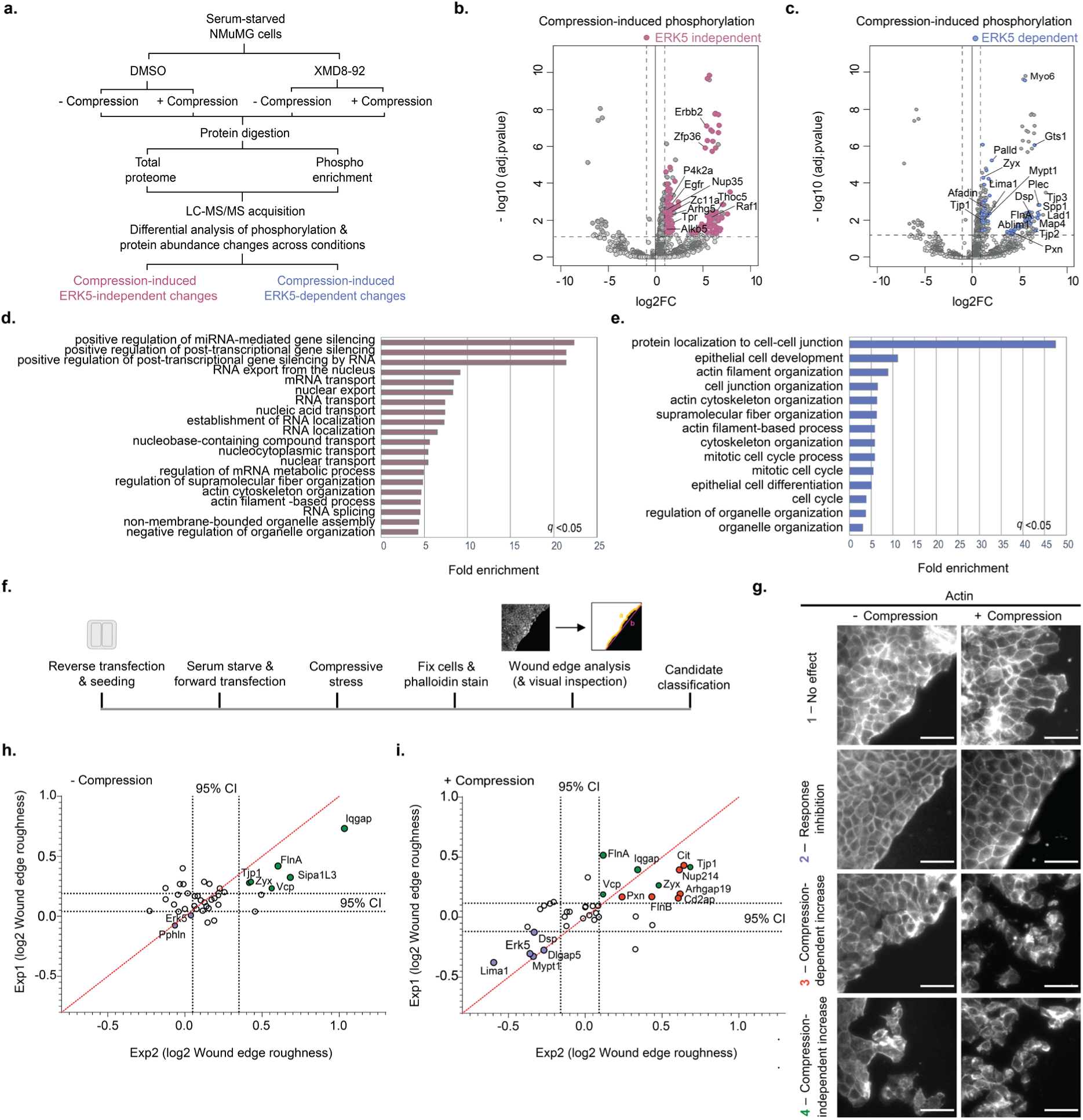
Phospho-proteomic and siRNA-based screening identified ERK5 dependent signaling candidates regulating the actin cytoskeleton and cell-cell junctions. **a.** Experimental workflow used for phospho-proteomic profiling of NMuMG cells to classify compression-regulated, ERK5-dependent and independent phospho-sites. Phospho-peptides were identified by MS-analysis after F_e_-NTA enrichment, using extracts prepared from serum-starved NMuMG cells exposed (filled circles) or not (open circles) to compression for 3 hours, with or without ERK5 inhibition by XMD8-92. **b.** - **e.** Volcano plot showing differential protein phosphorylation in compression versus no-compression comparison (p-adj. ≤ 0.05, log2FC>1.0) marking ERK5-independent (**b**) and ERK5-dependent (**c**) compression-induced phosphosites. Proteins involved in cell-to-cell contact and the regulation of the actin cytoskeleton (**c**) and gene expression regulation and nuclear import (**b**) are highlighted, respectively. Each phosphorylated protein is represented by its most significant phosphopeptide. The p-value <0.05 is considered significant and corresponds to an ANOVA two-sided p-value, adjusted for multiple testing across the cells-treatment conditions of interest after post hoc analysis (FDR). Gene ontology (GO) enrichment analysis revealed ERK5-dependent targets related to cell-cell junction and actin cytoskeleton organization (**e**), and pressure-induced, but ERK5-independent targets related to nuclear transport regulation and post-transcriptional gene silencing (**d**). Bonferroni- adjusted p-value < 0.05 corresponds to two-sided Fisher’s exact test. All genes in Mus musculus database are used as reference list. **f.** Experimental workflow to functionally validate candidate ERK5-substrates. Candidates were siRNA depleted in serum starved NMuMG cells growing in ibidi chambers and examined by the epithelial edge assay after 24 hours of compression. The screen was repeated twice, each time dividing the candidates into five or six batches, with each batch including non-targeting siRNAs and siERK5 controls. Cells were fixed and stained with rhodamine phalloidin and the edge roughness imaged and quantified. The resulting phenotypes were grouped in 4 functional categories, which are illustrated by the edge images of NMuMG cells exposed (+) or not (-) to compressive pressure **g**. Category 1: siRNA knockdown has no influence on the phenotypic response to compression; category 2: siRNA knockdown prevents the response to compression; category 3: siRNA knockdown increases cell dissociation upon compression; category 4: siRNA knockdown leads to cell dissociation even in the absence of compression. **h.** and **i.** Graphic comparison of wound edge roughness of two screen replicates. The wound edge roughness of each target is normalized to the average value of the control siRNAs and log2 transformed. The dashed lines indicate 95% confidence intervals (95% CI) around the median of all measurements in each independent screen. Candidates that significantly reduce the wound edge perimeter upon compression compared to cells treated with control siRNAs are marked in blue. Candidates enhancing the wound edge roughness with (**i**) or without compression (**h**) are marked in red and green, respectively.

### siRNA-based screen identified signaling candidates important for invasive response to compression

To functionally validate relevant ERK5-substrates, we selected a total of 37 candidates (Suppl. Tables Fig. 4) from the phosphoproteomic analysis associated with actin regulation or cell-cell junctions. These candidates were then depleted by siRNA, and the cellular phenotype analyzed with or without compression for 24 hours using the tissue edge assay (Fig. 4f). Based on this analysis, we distinguished four phenotypic categories (Fig. 4g). Category 1 includes genes for which siRNA depletion did not change the edge roughness compared to controls. Among these candidates is the transcription factor Yes-associated protein-1 (YAP1), which is dispensable for rapid morphological changes but may be important for fine-tuning and/or long-term adaptation of cells responding to compressive mechanical stress. Category 2 comprises genes that phenocopy ERK5 depletion, e.g. their absence prevents the invasive response of NMuMG cells to mechanical compression (Fig. 4g, i). We hypothesize that ERK5-dependent phosphorylation of these proteins is required to establish the cytoskeletal and morphological changes of cells exposed to compressive stress. Indeed, proteins of this group, such as Lima1 and Myosin phosphatase target subunit 1 (MYPT1), have previously been found to regulate actin bundling and contractility, respectively. In contrast, depleting candidates clustered in category 3 display the opposite effect – they enhance the invasive response upon compression. Proteins in this category include the junction-associated proteins ZO1 and ZO2 and actin-binding molecules regulating actin filament organization such as CD2-associated protein (Cd2ap), Zyxin, Filamin A, and Filamin B. Cell-cell junctions in NMuMG cells lacking these candidates appear normal, but cell-cell detachment and edge roughness significantly alters upon compression (Fig. 4g, i). Many of these candidates stabilize cell-cell contacts, and ERK5 phosphorylation may thus impair their protective function, thereby enhancing cell-cell detachment. Surprisingly, siRNA depletion of the nuclear pore complex subunit Nup214 augmented cell-cell detachment, and compression also caused sickle-shaped nuclear morphology. In contrast, siRNA depletion of Paxilin increased edge roughness not by enhancing cell-cell separation but rather through increased protrusions upon compression. Finally, category 4 is composed of candidates for whom depletion increases cell-cell detachment and edge roughness even in the absence of compression (Fig. 4g, h). Some of these candidates overlap with category 3, highlighting their importance in preserving robust cell-cell junctions and epithelial morphology. Thus, we hypothesize that ERK5 targets in categories 3 and 4 may increase ERK5 activity or more likely, stabilize cell-cell junctions. If so, their ERK5 dependent phosphorylation may help to dissolve cell-cell junctions upon compression.

### ERK5 phosphorylates MYPT1 on Ser909, thereby decreasing peripheral tension

We decided to substantiate the role of ERK5-dependent phosphorylation of the category 2 gene MYPT1, a subunit of the myosin light chain phosphatase (MLCP) responsible for its targeting to myosin^29^. Active MLCP dephosphorylates and in turn inactivates myosin light chain (MLC), shifting the balance towards reduced myosin activity (Fig. 5a), thereby decreasing actin contraction and cortical tension^29,30^. We investigated the effect of compressive stress on MLC activity by comparing ERK5-dependent MLC phosphorylation in the presence and absence of compression (Fig. 5b). Compressed NMuMG cells were immunostained with phospho-MLC specific antibodies, and for control, RNAi-depleted of ERK5 or MYPT1 (Fig. 5b). Interestingly, mechanical compression significantly reduced phospho-MLC levels at the cell periphery compared to uncompressed cells (Fig. 5b, c). In contrast, compressed cells lacking ERK5 or MYPT1 maintained phospho-MLC and preserved its peripheral localization. Cytoplasmic and nuclear localization of total MLC did not change during compression or in the absence of ERK5 and MYPT1 (Extended Data Fig. 6a). In addition, non-muscle myosin IIA remained localized at cell-cell contacts after 3 h of compression (Extended Data Fig. 6b), indicating that the loss of active myosin is due to dephosphorylation rather than the loss of cell contacts. To validate that ERK5-MYPT1 signaling regulates actomyosin tension at cell-cell adhesions we stained NMuMG cells for vinculin, a mechanosensitive protein recruited to the cadherin-catenin complex under tension^31^. Indeed, compression led to a reduced vinculin localization at the cell periphery in an ERK5- and MYPT1-dependent manner (Extended Data Fig 6c). We thus conclude that ERK5 signaling inhibits myosin activity to disrupt tension at cell-cell contacts, which promotes invasive morphological changes upon compression.

**Fig. 5.**
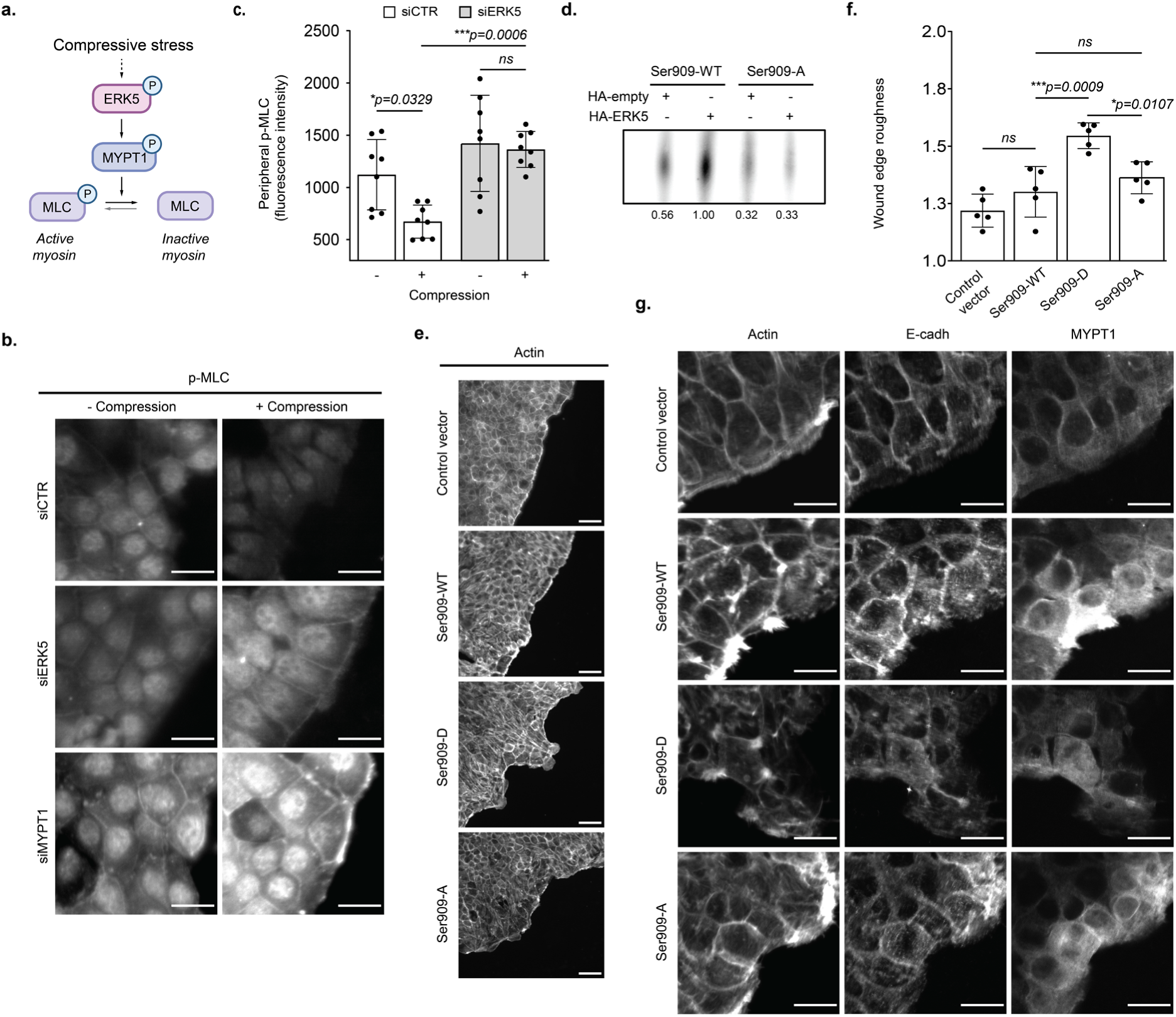
ERK5 phosphorylates MYPT1 on Ser909, thereby decreasing peripheral tension. **a.** Schematic illustration of the ERK5 signaling pathway phosphorylating S909 of the myosin light chain phosphatase subunit MYPT1, which in turn dephosphorylates and thereby activates the myosin light chain (MLC). Active myosin controls cortical tension by pulling on actin fibers. **b**. and **c**. NMuMG cells siRNA depleted for ERK5 (siERK5), MYPT1 (siMYPT1) or treated with a control oligo (siCTR) were exposed (+) or not (-) to physical compression for 3 hours, and immunostained for phospho-MLC (p-MLC). Note that MLC phosphorylation is decreased upon compression by an ERK5 and MYPT1-dependent mechanism. Scale bar: 20 μm (**b**). Fluorescence intensity of p-MLC at cell edges was quantified as described for phospho-ERK5. Compression significantly reduces epithelial p-MLC levels while ERK5 depletion protects p-MLC at cell-cell contacts (**c**). **d.** Extracts prepared from cells expressing HA-tagged ERK5 (HA-ERK5) or carrying an empty control plasmid (HA-empty) were incubated with HA-affinity resins. Immobilized proteins were incubated with a peptide encompassing the S909 phosphorylation site of MYPT1 (S909-WT) or a mutant peptide where Ser909 was replaced by an alanine residue (S909-A) in the presence of γ-^32^P-ATP. The peptides were separated by thin layer chromatography (TLC) and analyzed by autoradiography. The numbers below represent the densitometric quantification of phosphorylated peptides relative to active ERK5 upon compression. **e. - g.** NMuMG cells grown in ibidi chambers were reverse-transfected with an empty control vector or a vector overexpressing wild-type MYPT1 (Ser909-WT), or MYPT1 mutants changing S909 either to a phosphomimicking glutamic acid (Ser909-D) or a non-phosphorylatable alanine (Ser909-A) residue. Wound edge roughness was visualized by phalloidin staining (**e**) and quantified to compare the different conditions (**f**). N=5; ANOVA with Tukey multiple comparison; ***p<0.001; p<0.05, considered statistically significant. Note the elongated morphology and significantly weaker intercellular connections in NMuMG cells expressing Ser909-D compared to Ser909-WT and Ser909-A mutants (**g**).

Interestingly, the phosphoproteomic analysis identified serine 909 (Ser909) in MYPT1 as a compression-regulated, ERK5-dependent phosphorylation site. To corroborate these data, we performed targeted quantification of Ser909 phosphorylation by parallel reaction monitoring (PRM). Indeed, Ser909 phosphorylation was reduced when ERK5 activity was inhibited with XMD8-92 in compressed NMuMG cells, while MYPT1 protein levels remained unaffected (Extended Data Fig. 6d, e). To examine whether ERK5 directly phosphorylates MYPT1, we immobilized HA-ERK5 on HA-coated beads and incubated the affinity purified kinase (HA-Erk5) with a MYPT1 peptide encompassing S909 (Ser909-WT) in the presence of γ^33^P-ATP. Indeed, thin layer chromatography (TLC) confirmed *in vitro* phosphorylation of the peptide (Fig. 5d, Extended Data Fig. 6f), which was absent in untagged controls (HA-empty) or when using a control peptide where the phosphorylated serine residue was replaced by a non-phosphorylatable alanine (Ser909-A). Together, these data suggest that ERK5 directly phosphorylates MYPT1 on S909 upon mechanical compression.

We next tested whether MYPT1-S909 phosphorylation is functionally relevant for the invasive phenotype triggered by compression. To this end, we measured edge roughness in NMuMG cells overexpressing MYPT1 mutants in which Ser909 was substituted with aspartic acid (Ser909-D) to mimic phosphorylation or alanine (Ser909-A) to prevent ERK5 phosphorylation. Indeed, cells overexpressing Ser909-D showed significantly increased edge roughness even in the absence of compressive stress (Fig. 5e, f), which was not observed with a control vector or when overexpressing wild-type MYPT1 (Ser909-WT) or the non-phosphorylatable Ser909-A mutant (Fig. 5e, f). Interestingly, cells overexpressing Ser909-D exhibit an elongated morphology, reduced peripheral junctional actin and significantly weaker intercellular E-cadherin connections (Fig. 5g). This dominant phenotype was not a consequence of higher expression of Ser909-D compared to Ser909-WT or Ser909-A (Fig. 5g, Extended Data Fig. 6g). Taken together, these functional assays suggest that MYPT1 phosphorylation by ERK5 is necessary and sufficient to promote the invasiveness of NMuMG cells upon mechanical compression.

### ERK5 and ROCK antagonistically regulate MYPT1 and reduce peripheral tension upon compression

The Rho associated protein kinase ROCK is well known to phosphorylate MYPT1 on Thr694 and Thr852 (Thr696 and Thr853 in humans), the latter site being ROCK-specific^32^. Phosphorylation of Thr852 decreases MYPT1 binding to myosin, thus promoting myosin phosphorylation^33^. As expected, immunofluorescence staining of NMuMG cells with antibodies specifically recognizing phospho-Thr852 was diminished when the ROCK kinase was depleted by siRNA (Fig. 6a). Interestingly, phospho-Thr852 was also reduced after 3 h of mechanical compression independent of ERK5 (Fig. 6a), suggesting that compressive stress inhibits ROCK and in turn promotes activity of the myosin light chain phosphatase (MLCP). To test whether ROCK and ERK5 signaling might antagonistically regulate MYPT1, we performed tissue edge assays in NMuMG cells treated with the ROCK inhibitor Y27832, with or without simultaneous inhibition of ERK5 by XMD8-92. As expected, inhibition of ROCK significantly increased cell-cell detachment enhancing edge roughness upon compression (Fig. 6b, c). However, inhibition of both ROCK and ERK5 activities restored the smooth edge and prevented the invasive phenotype (Fig. 6b, c), implying that ROCK and ERK5 antagonize each other. We conclude that compressive mechanical stress decreases peripheral tension by two independent mechanisms: compressive stress inactivates ROCK, thereby reducing Thr852 phosphorylation of MYPT1 which promotes MLCP activity. In addition, compressive stress activates ERK5, which phosphorylates MYPT1 on Ser909, thereby fully activating MLCP. As a result, these synergistic mechanisms lead to rapid de-phosphorylation of MLC, which in turn enhances an invasive phenotype by reducing peripheral tension and promoting dissolution of cell-cell contacts (Fig. 6d).

**Fig. 6.**
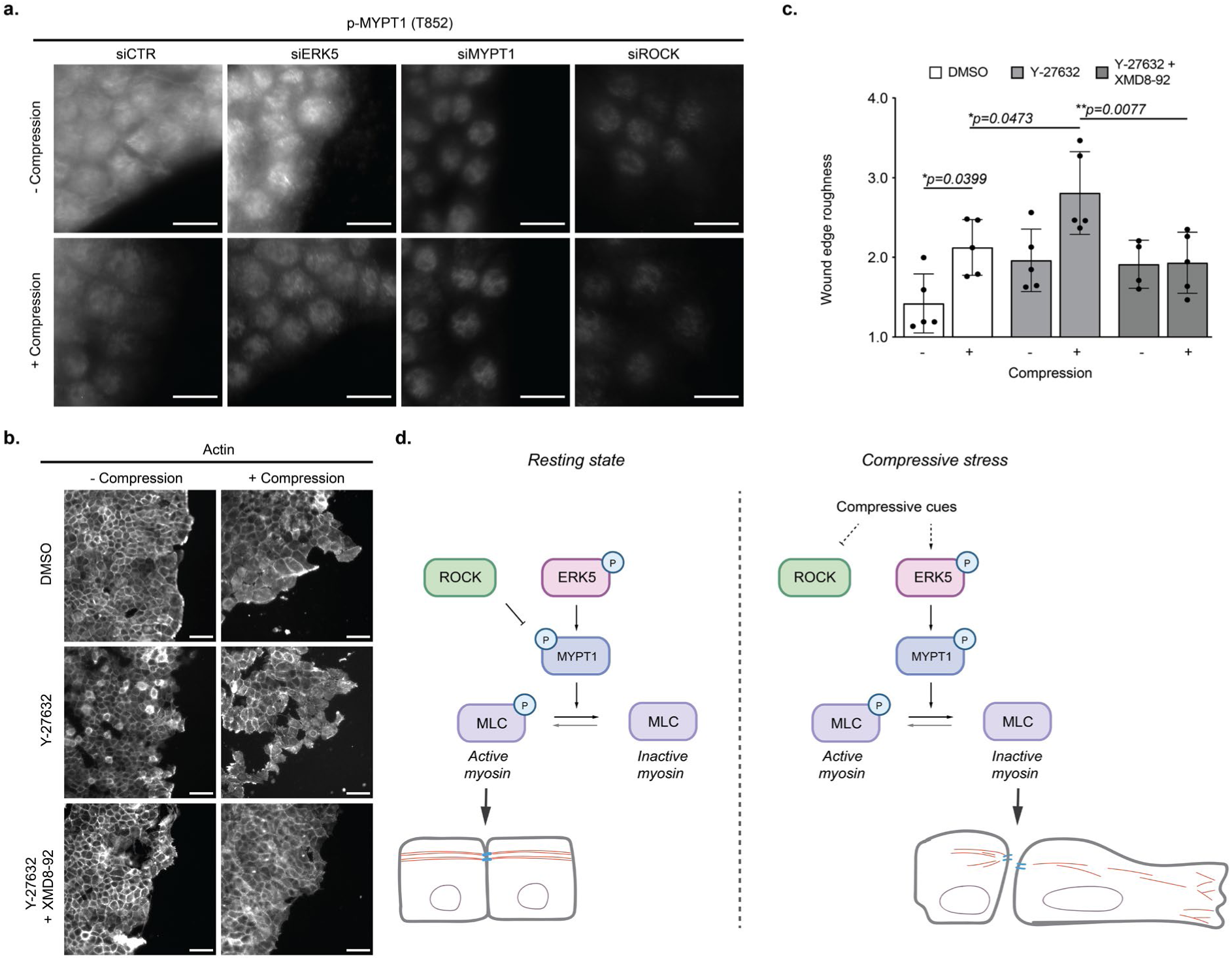
ERK5 and ROCK cooperatively regulate MLC upon compressive stress, thereby promoting cell invasion. **a.** NMuMG cells siRNA depleted for ROCK (siROCK), ERK5 (siERK5), or both together (siROCK + siERK5), or treated with a control oligo (siCTR), were exposed (+) or not (-) to physical compression for 3 hours, and phospho-threonine 852 (T852) of MYPT1 (p-MYPT1) was analyzed by immunofluorescent staining. Scale bar: 20 μm. Note that the ROCK-dependent T852 phosphorylation is decreased upon compression. **b.** and **c.** NMuMG cells grown in ibidi chambers were treated with the ROCK inhibitor Y27632, its solvent DMSO, or together with the ERK5 inhibitor XMD8-92. Cells were exposed (+) or not (-) to compressive pressure for 24 hours, and wound edge roughness was quantified after phalloidin staining. Note that ROCK and ERK5 activities antagonize each other. N=5; ANOVA followed by post hoc Bonferroni multiple comparison of relevant conditions; ***p<0.001; p<0.05 considered statistically significant. **d**. During unstressed conditions, ROCK phosphorylates MYPT1 on its inhibitory Thr852 site (Thr853 in human), thereby increasing active myosin (phospho-MLC) and stabilizing actin structures and cell-cell junctions. Compressive stress inhibits ROCK but activates ERK5, which then phosphorylates MYPT1 on Ser909. This phosphorylation switch activates the myosin light chain phosphatase complex (MLCP), leading to myosin inhibition and in turn changes in cortical actin structures and disruption of cell-cell junctions, thus promoting an invasive behavior.

## Discussion

Our study demonstrates that compression is sufficient to trigger invasive features of untransformed epithelial cells grown in 2D, and similarly, compression increases leader cell formation in pancreatic and breast cancer cell lines^6,9^. Compressive stress disrupts mechanical coupling of cells through adherent junctions and the protective barrier function of tight junctions, thereby promoting migratory features.

Several lines of evidence suggest that the MAP kinase ERK5 plays a key role in signaling cellular invasion upon compressive stress. First, compressive stress activates ERK5 at the cell periphery, and ERK5 also responds to shear stress^15,16^ and cytokines^34^. How compressive stress is sensed and transduced intracellularly to the MEK5-ERK5 module remains to be investigated. This could either occur at the plasma membrane, for example, by an integrin-dependent mechanism^1,35^ or by engaging mechano-sensors in the nuclear envelope such as the LINC complex^36^. Indeed, compressive stress induces nuclear deformation in our device, and this phenotype is independent of ERK5 activity. Second, active ERK5 is necessary and sufficient to rearrange the actin cytoskeleton and disassemble cell-cell contacts. Consistent with this notion, we identified multiple ERK5 substrates involved in adherence junctions and cytoskeletal dynamics, and these targets were not previously identified as ERK5 effectors. Third, mechanistically, ERK5 directly phosphorylates serine 909 of MYPT1, which in turn dephosphorylates MLC, thereby relaxing peripheral actomyosin tension necessary to maintain cell-cell contacts. Strikingly, NMuMG cells expressing a phospho-mimicking S909D mutant exhibit many invasive features such as weakened adherens junctions and peripheral actin reorganization, implying that S909 phosphorylation comprises a key event for compressive stress signaling (Fig. 5e, g). In unstressed cells, MYPT1 is a well characterized substrate of the ROCK kinase, which inhibits MLCP activity by phosphorylating T852 and T696^29,32^. Interestingly, compressive stress inhibits ROCK activity, and ERK5 and ROCK antagonize each other to cooperatively increase MLCP activity, in part because S909 phosphorylation may sterically interfere with ROCK binding^37^. How phosphorylation of S909 by ERK5 stimulates the activity of MLCP remains unclear, but it does not involve increased expression levels. Additional compressive stress responsive ERK5 substrates were identified during our studies, and future investigations will pursue physiological and mechanistic underpinnings of the coordinated ERK5 signaling response.

Cells dynamically regulate actomyosin tension to modulate cell shape and intercellular junction strength^31,38^. The integrity of cell-cell junctions is regulated through a tight balance of myosin activity^38^. Active junctional myosin sustained by RhoA-ROCK activity is indispensable for mechanically coupling cells through adherens junctions and maintaining their integrity, while exacerbated activity can break contacts apart^30–31,38–39^. We find that in response to compressive stress ERK5 is required for cells to decrease cortical actomyosin tension thus weakening cell-cell contacts, promoting an invasive phenotype. Thus, decreased ROCK activity and ERK5 activation cooperate to rapidly dissociate cell-cell contacts upon compression. Interestingly, ERK1/2 activity waves are associated with increased actomyosin contraction during collective cell migration^40^, raising important questions about spatiotemporal differences among distinct MAP-kinase modules during mechanical compression.

While activation of the MEK5-ERK5 module provides an important signaling axis regulating cell-cell interactions and cytoskeleton dynamics, we also discovered proteins that are phosphorylated upon compression by an ERK5-independent mechanism. These proteins are enriched in nuclear factors involved in gene silencing, RNA biology, nuclear transport, or chromatin organization, suggesting that compressive stress may trigger nuclear re-organization. The kinase(s) mediating these effects are unknown but may include ATM/ATR, which prevents nuclear rupture during DNA replication and repair^41^. Moreover, nuclear deformation may promote nuclear transport and activation of transcription factors to orchestrate changes in gene expression^42^. Further work will be needed to understand the physiological and pathological implications of these ERK5-independent pathways in the compressive stress response.

Taken together, our findings reveal the cellular signaling network activated by compressive mechanical stress and identify the MEK5/ERK5 MAP kinase module as a key regulator of cell-cell contacts and cytoskeletal dynamics. ERK5 is often upregulated and overactive in cancer cells^19–22^ and required for EMT^17^, a critical process in cancer progression, promoting cell migration and invasive traits^18^. Given the rapidly emerging relevance of ERK5 for tumor progression and metastasis, understanding its molecular function and upstream regulators may reveal new therapeutic strategies.

## Author contributions

Conceptualization, N.S., FvD, and M.P.; formal analysis, N.S., FvD, and MP; funding acquisition, M.P.; proteomics, N.S., T.S. and F.U.; microscopy, image analysis and quantifications, N.S., and FvD., together with ScopeM; *in vitro* kinase assays, FvD.; Visualization, N.S., T.S., and F.U.; Supervision, M.P., and FvD; writing – original draft, N.S. and M.P.; writing – review and editing, N.S., FvD and M.P., with inputs from Alicia Smith and all authors.

## Acknowledgment

We thank the ETH Scientific Center for Optical and Electron Microscopy (ScopeM) and in particular Dr. Tobias Schwarz for assistance with microscopy, and the ScopeM Image clinic and Zurich Image and Data Analysis School (ZIDAS) for help with image quantifications. We are grateful to Gerhard Christofori for the NMuMG cell line, Daniel Smith for fabricating the loads, Ranjan Mishra for initial observations, and Teodora Urosevic, Olena Marchenko and Anna Huyen Hgo for experimental support during their laboratory internships. We thank Gabriel Neurohr and members of the Peter lab for discussions and critical comments on the project, and Alicia Smith for editing. This work was supported by the Swiss National Science Foundation (SNSF) and ETH Zürich.

## Methods

### Cell culture

Murine mammary gland epithelial cell line NMuMG, was obtained from the laboratory of G. Christofori and originally from American Type Culture Collection (ATCC). Cells were cultured in DMEM (gibco) supplemented with 10% FBS and 1% penicillin/streptomycin at 37°C under 5% CO_2_.

### Plasmid construction

HA tagged dominant active MEK5 (pcDNA3-MEK5DD-HA) was purchased from Addgene (Addgene plasmid 65247). To generate the *pMEF2-GFP* reporter a 3XMEF2-luc reporter plasmid (Addgene plasmid 32967) was utilized to replace luciferase by GFP using Nco1/Sac1 restriction sites. MYPT1 S909D and S909A mutations were introduced by round-the-horn (whole plasmid) PCR strategy on pK.myc-WT-MYPT1 (Addgene plasmid 24101).

### Compressive stress application

An in-house built metal load of a specified weight applies constant force to an agarose disc that is in contact with cells grown on a transwell membrane of 0.4 µm (Corning® 24 mm Transwell® 3450, Falcon® Permeable Support 353090) or 8.0 µm pores (Corning® 24 mm Transwell® 3428). NMuMG cells were washed once with PBS, trypsinized, resuspended in culturing medium and seeded at a desired density onto the transwell membrane with transwell inserts residing in a 6-well plate. The medium is placed below the transwell insert in a 6-well plate enabling nutrient and oxygen exchange between the upper and lower compartment. After 24 h of growth, the culturing medium was aspirated, cells washed with sterile PBS once and serum-deprived medium (DMEM + 1% penicillin/streptomycin) placed within the upper and lower compartment. After 24 h of serum starvation, the agarose cushion is placed on top of the cells, while compression-applying loads are placed onto the agarose cushion. The weight of one load is 24.5 g with a diameter of 24 mm, which translates to mechanical compression of 4 mmHg. For the control samples where no compression is applied only the agarose cushion was placed on top of the cells. Agarose cushion was prepared from 1% agarose in PBS, which was briefly microwaved to dissolve agarose. Subsequently the agarose gel of 1 mm thickness was made using vertical electrophoresis gel-casting system previously sterilized by washing in 70% ethanol and 1 hour incubation under UV light. The pressure-applying loads were sterilized by autoclaving.

### Drug treatment, siRNA transfection and construct overexpression

To inhibit ERK5 and ROCK, NMuMG cells were treated with 10 µM XMD8-92 (Tocris Bioscience, 1234480-50-2) or Y-27632 (Sigma-Aldrich, Y0503), respectively, during serum starvation (24 h prior to compressive stress application) and at the time of compressive cue application. Control cells without ERK5 inhibition were treated with DMSO. For siRNA-mediated silencing cells were reverse transfected with Lipofectamine RNAiMax reagent (Thermo Fisher) and 40 nM siRNA/s (2:1 ratio) at the time of seeding. During serum starvation, cells were subjected to forward transfection with Lipofectamine RNAiMax reagent and 40 nM siRNA/s. To overexpress constructs, NMuMG cells were reverse transfected at the time of seeding with 40 ng of plasmid DNA per ibidi well or 840 ng of plasmid DNA per transwell surface using Polyplus jetOPTIMUS DNA transfection reagent (0.04 µL reagent in 15 µL total transfection mixture volume per ibidi well or 0.84 µL reagent in 250 µL total transfection mixture volume per whole transwell insert).

### Transmigration assay

The transmigration assay was performed using a transwell-based system for compressive stress application with 8.0 µm pore polyester membrane. NMuMG cells were seeded at the density of 300 000 cells/insert in the upper compartment in 1.5 mL total volume of culturing medium, while 2.5 mL of the culturing medium was placed below the insert in the lower compartment. The cells were grown for 24 h at 37°C in an atmosphere containing 5% CO_2_. Next, cells were serum starved for 24 hours. To silence ERK5 activity, cells were additionally treated with XMD8-92 or transfected with siRNAs against ERK5, as described. After 24 hours of compressive force application, the transwell inserts were discarded while the lower compartment (6-well plate) was replenished with 10% FBS and left at 37°C and under 5% CO_2_ atmosphere for 24 h for transmigrated cells to adhere and form colonies. Colonies were fixed with 4% PFA for 15 min, washed with PBS once and incubated with 0.25% crystal violet (w/v) for 30 min. Colonies were washed with PBS until clear. Plates were scanned and the colonies manually counted in ImageJ^43^.

### ibidi wound-edge roughness assay

Cells (2×10^4^/well) were seeded in Culture inserts 2 well (Ibidi, 80209) placed onto the 0.4 µm pore polyester membranes of transwells in compressive stress platform. After 24 h of attachment and growth the medium was exchanged to serum-deprived medium for 24 h within the Culture inserts and lower compartment. Next, culture inserts were gently removed by tweezers leaving a cell-free gap and compressive stress was applied for 24 h. The cells were fixed on the transwell membrane with 4% PFA for 15 min. Cells were washed once with PBS and the membrane piece with two cell layers and a gap was cut and stained with rhodamine phalloidin (Invitrogen, R415) and 1 µg/mL DAPI (Invitrogen, 62248). The cells were mounted on microscopy slides, sealed with cover glass (Epredia, 24×40 mm) and imaged with Nikon-Ti-Eclipse system using 20x or 10x objective (Plan Apo 20x NA 0.45, Plan Apo 10x NA 0.45) using Micro-Manager (version 5.02.00) software^43^. Minimum of five images of the edges of two cell layers (wound edges) were acquired per condition and subjected to further analysis in ImageJ^44^.

### Immunofluorescence and microscopy

NMuMG cells were grown and serum starved as described for the ibidi wound-edge roughness assay and either reverse transfected with plasmids or siRNA/s at the time of seeding or treated with the inhibitors as described. Cells were subjected to compression for 3 h or 24 h as described in figures. After compressive stress application, cells were then washed with PBS and fixed with 4% paraformaldehyde in PBS for 15 min. Cells were washed with PBS and blocked with 2% bovine serum albumin (BSA) 0.1% Tween-20 in TBS. Membrane parts with two cell layers were excised and incubated with the primary antibodies in blocking buffer (2% BSA 0.1% Tween-20 in TBS) at +4°C overnight in a humid chamber. The cells were then washed three times with blocking buffer (10 min each wash), stained with secondary antibodies (1:100), rhodamine phalloidin and DAPI in blocking buffer for 2 h and finally washed three times with PBS (10 min each wash). The antibodies are provided in the Extended Data Table 1. The cells were mounted on the microscopy slides using Vectashield mounting reagent and sealed with the cover glass (Epredia, 24×40 mm). Images were recorded by Zeiss LSM880 confocal microscope system using 63x objective (DIC Plan-Apochromat 63x NA 1.4 oil) with ZEN Blue software and Nikon-Ti-Eclipse system using 60x objective (Plan Apo 60x NA 1.4 oil) with Micro-Manager (version 5.02.00) software^43^. Immunofluorescence images of phospho-threonine 852 (T852) of MYPT1 were acquired with Nikon-Ti2-Eclipse using 60x objective (Plan Apo 60x NA 1.4 oil) and NIS Elements AR 5.42.03.

### Image analysis

#### Wound-edge roughness

To analyze the wound edges a custom-developed script, written in the ImageJ macro language, was employed. Briefly, the wound edge images blurred, thresholded and converted to mask, upon which the perimeter is obtained. To obtain the only perimeter of the wound the perimeters of individual image edges are subtracted. To compute the wound edge roughness index the wound perimeter is normalized to the length of the straight line connecting two edge points of the wound. The macro is applied in batch-processing mode of ImageJ.

#### Cell and nuclear morphology

DAPI images of cells at the wound edge acquired with Nikon-Ti-Eclipse 60x objective (Plan Apo 60x NA 1.4 oil) and Micro-Manager software^43^ were segmented with Cellpose (2.0)^45^ wrapper in Fiji using the pre-trained nuclei model. To quantify cell shape, cells imaged at the wound edge with Nikon-Ti-Eclipse 60x objective (Plan Apo 60x NA 1.4 oil) were segmented with Cellpose (2.0)^45^ wrapper in Fiji using the pre-trained cell model and composite DAPI and rhodamine phalloidin images as an input. To measure nuclear and cell elongation, aspect ratio of segmented nuclei and cells was computed in Fiji, respectively. The steps were developed into ImageJ macros for image batch processing. The aspect ratios were measured on more than 300 cells per condition.

#### p-ERK5 and p-MLC fluorescence intensity

Cells were segmented based on composite images of DAPI and rhodamine phalloidin-stained cells with Cellpose (2.0)^45^ in Fiji using the pre-trained cell model. To quantify fluorescence intensities, a Fiji script was customized. The script imports Cellpose cell segmentations saved as regions of interest (ROIs) and performs enlargement of each ROI by a few pixels to ensure the total cell intensities are measured, followed by band creation around the cell periphery to measure fluorescence intensity at the cell edge. For each position, total and peripheral fluorescence intensities are measured using maximum intensity projections of 10 z-planes using DAPI channel as a reference. The intensities were measured for approximately 300 cells at the wound margin per condition.

### Western blot

Cells were grown on a transwell platform at 300 000 cells/well density with three transwells seeded per condition. After treatments as described in figures, transwells were flash frozen in liquid nitrogen and the cells were lyzed in 200 µL/well Laemlli buffer by scraping on ice. Samples were heated at 70°C for 5 min, separated by SDS-PAGE and transferred onto PVDF membrane (Immobilon^R^-P Transfer Membrane) using semi-dry system (Hoefer^TM^ TE 77). Membranes were blocked for 1 hour in 2% BSA 0.1% Tween-20 in TBS, followed by overnight incubation with primary antibodies diluted in blocking solution. Membranes were washed three times in TBST, followed by incubation with a HRP-linked goat anti-rabbit or anti-mouse IgG secondary antibody (1:2,000, Cell Signaling Technology) for chemiluminescence detection. The list of primary and secondary antibodies is provided in Extended Data Table 1. The signal was developed using SuperSignal™ West Pico PLUS or Femto Chemiluminescent Substrate solution (Thermo Fischer) and scanned on Fusion FX7 imaging system (Witec AG).

### Phosphoproteomics

#### Mass spectrometry sample preparation

For analysis of compression- and ERK5-dependent phosphorylation events NMuMG cells were seeded on a transwell platform in three transwells per replicate at a density of 300 000 cells/well. Each condition was prepared in triplicate. The cells were grown, serum starved and pre-treated with 10 µM XMD8-92 as described. On the day of compression application, serum-deprived medium with DMSO control or 10 µM XMD8-92 was re-added and compressive stress applied for 3 h. The cells were lysed by scraping on ice in 200 µL/well of urea lysis buffer containing 8 M urea, 50 mM Tris pH 7.5, 75 mM NaCl, 10 mM NaF, 10 mM pyrophosphate, 5 mM β-glycerophosphate and cOmplete^™^ Protease Inhibitor Cocktail tablets (Roche). The lysates of the same sample replicate were pulled and subjected to five founds of 30 s sonication at 80% amplitude with 30 s breaks on ice. The samples were centrifuged for 20 min at room temperature and the supernatants transferred to fresh tubes on ice. The protein concentrations were measured with BCA assay following manufacturer’s instructions (Thermo Scientific™ Pierce™ BCA Protein Assay Kit). After correcting the samples for protein concentration, reduction was performed with 5 mM tris-2-carboxyethyl-phosphine (TCEP) at 700 rpm agitation and 37°C for 30 min, followed by alkylation with 10 mM iodoacetamide at 700 rpm and 37°C for 30 min. The samples were diluted two times in 50 mM ammonium-bicarbonate (AMBIC) to reach 4 M concentration of urea. First round of proteolysis was done with 1:150 enzyme/sample protein of LysC for 1 h at 700 rpm agitation and 37°C. The concentration of urea was reduced to 2 M with 50 mM AMBK and second proteolysis step done with 1:50 enzyme/sample protein of trypsin overnight at 700 rpm agitation and 37°C. Proteolysis was quenched with 5% trifluoroacetic acid (TFA) and the samples were centrifuged for 10 min at maximal speed before C18 cleanup following manufacture’s instruction. The samples were dried and re-suspended in 20 µL of 70% acetonitrile (ACN) 0.9% TFA followed by sonication in the water batch for 10 min. For the full proteome analysis 10% of the sample was diluted in 50 µL of 2% ACN. The rest of the sample was diluted in 200 µL of binding buffer for phosphopeptide enrichment. Phosphopeptides were enriched using High-Select™ Fe-NTA Phosphopeptide Enrichment Kit (Thermo Scientific) following manufacture’s instruction. Phosphopeptides were desalted using C18 Ultra Micro Spin Columns, dried and re-suspended in 0.1% formic acid (FA).

#### Mass spectrometry acquisitions

Untargeted experiment: Samples were acquired in DDA mode in an Orbitrap QE Exactive Plus (Thermo) equipped with a nanoelectrospray ion source coupled to EASY-nLC-1000 LC system (Thermo). Peptides were separated using a reverse phase column (75 µm ID x 400 mm New Objective, in-house packed with ReproSil Gold 120 C18, 1.9 µm, Dr. Maisch GmbH) in a gradient of 180min from 3 to 25 (160min) and 25 to 40% (20min). The composition of MS is buffer A: 0.1% (v/v) formic acid; buffer B: 0.1% (v/v) formic acid, 95% (v/v) acetonitrile. The acquisition was set to perform one MS1 scan followed by a 20 MS2 scans for top 20 ion peaks; MS1 scans (R=70’000 at 400 m/z, AGC=3e6 and maximum IT= 64ms), HCD fragmentation (NCE=25%), isolation windows (1.4m/z) and MS2 scans (R=35’000 at 400 m/z, AGC=1e5 and maximum IT=110ms). A dynamic exclusion of 30s was applied, and charge states lower than two and higher than six were rejected for the isolation. The dataset, including raw data and output from MaxQuant research are available in the PRIDE repository.

Targeted analysis was performed on an Orbitrap Exploris480 mass spectrometer (Thermo Fisher) coupled to an Vanquish Neo liquid chromatography system (Thermo Fisher). Peptides were separated using a reverse phase column (75 μm ID x 400 mm New Objective, in-house packed with ReproSil Gold 120 C18, 1.9 μm, Dr. Maisch GmbH) across a gradient from 7 to 50% in 60 minutes (buffer A: 0.1% (v/v) formic acid; buffer B: 0.1% (v/v) formic acid, 80% (v/v) acetonitrile). A targeted scheduled method was set up with one MS1 scan (R=120’000 at 400 m/z, normalized AGC = 1e6 and max. IT auto) and targeted scans (R= 15’000, AGC = 1e5 and max. IT 100ms) using an isolation window of 1.8m/z and normalized collision energy 30%. The dataset, including raw data and output from Skyline analysis are available in the PRIDE repository.

#### Mass spectrometry data analysis

Acquired spectra in the untargeted analysis were searched using the MaxQuant software^46^ package version 1.5.2.8 embedded with the Andromeda search engine (PMID: 19029910) against Mus Musculus reference dataset (http://www.uniprot.org/, downloaded on 13.04.2020, 17’029 entries) extended with reverse decoy sequences. The search parameters were specific Trypsin, maximum two missed cleavage, carbamidomethyl as static peptide modification, Acetyl (Protein N-term); Oxidation (M), phosphorylation (S, T, Y) as variable modification and disabled “match between runs” option. The MS and MS/MS mass tolerance were set, respectively, to 20 ppm and 0.5 Da. False discovery rate of < 1% was used at the PSM level. Phosphorylation sites identified in the table “Phospho (STY)Sites.txt”. The normalized protein or peptide data matrices were exported from the MaxQuant software as CSV files for further quantitative analysis. In the absence of a confidently identified feature, the peptide and protein intensities were annotated as missing values (NA). To consider confidently identified single phosphosites (i.e., phosphorylation at S/T/Y) across experimental conditions, we first filtered the phosphopeptide datasets for those with at least two-thirds of the real experimental observations per three biological replicates of each condition (Supplementary Data). Missing values were imputed using an in-house script that sampled a random distribution of values, the mean of which corresponded to 95% of the minimal precursor peptide value (± 0.5 standard deviation (SD)) obtained from the respective phosphopeptide (or protein) measurements. Our filtered phosphoproteomic dataset contained less than 25% inputed values relative to the total number of quantified phosphopeptides (i.e., 6,964 phosphopeptides) and was used as the input file for further differential analysis. To monitor standard protein abundances in parallel with quantitative changes in phosphorylation, we also quantified protein abundances for the conditions of interest. We evaluated the data for statistical analysis using the Shapiro-Wilk normality test and generated histogram plots in R to observe the data distribution. We used a one-way ANOVA test followed by a Tukey’s HSD (honest significant difference) post hoc test to adjust the p-value for multiple testing across the respective cell-treatment conditions.

A differential analysis of phosphopeptides (or total proteins) was performed on four distinct conditions: (1) cells not subjected to mechanical compression (NP), (2) cells subjected to mechanical compression (P), (3) cells treated with XMD8-92 and not subjected to compression (NP Inh), and (4) cells treated with XMD8-92 and subjected to compression (P_Inh) (Fig. 4a). We made comparisons of interest to identify differentially modified and phosphorylated proteins under compressive stress by comparing phosphopeptides in relevant conditions (i.e., P vs. NP) and distinguishing sites that were phosphorylated by ERK5-dependent and - independent mechanisms (Supplementary Data, Figures 4a-c). We generated volcano plots from the log2-transformed fold change (FC) and the -log10-transformed p-values of proteins to reflect quantitative changes in phosphorylation under compressive stress (i.e., P vs. NP). To avoid redundancy in phosphoproteomic data visualization, each phosphorylated protein is represented by its most significant phosphopeptide (i.e., top phosphosite). Heatmaps were generated using the pheatmap R package (version 1.0.12) with scaled data and hierarchical clustering of log2-transformed relative signal intensities (total proteome or phosphosites) via Euclidean distances.

Functional enrichment analyses. For the protein-specific Gene Ontology (GO) biological processes (BP) annotation of the compression-induced phosphoproteome, we performed enrichment analyses relative to unique protein IDs (i.e., 348 unique protein IDs increased phosphorylation under compressive stress). Of the 348 phosphorylated proteins under compressive stress, 90 were ERK5-dependent and 258 were ERK5-independent. These were used as input lists in the PANTHER Overrepresentation Test (released October 13, 2022) via GO-enriched BP. A two-sided Fisher’s exact test was performed with reference list of all genes in Mus musculus database, and a Bonferroni-adjusted p-value (q-value) < 0.05 was considered significant. The top 20 biological processes (BP, q<0.05) were visualized using a bar plot and categorized as ERK5-dependent or ERK5-independent.

For targeted analysis, the peptide SAS[Pho]YSYLEDR (Mypt S909) was manually monitored using Skyline daily software^47^. Endogenous phosphorylated peptides were identified by matching the coelution and peak shapes of precursor and product ions with those of the reference heavy peptide (PEPotec Grade 3, Thermo). Quantification was performed the sum of the ten fragment ions per peptide. The intensity of endogenous peptides was normalized to the intensity of the heavy peptides, log2 transformed and the statistical analysis was performed with one-side unpaired t-test.

### *In vitro* TLC kinase assay

HA-ERK5 and empty vector containing HA were expressed in NMuMG cells and bound to HA-beads (Sigma) after lysis. Beads were incubated for 10 min at 37 °C with wild type (RLLGRSA**S**YSYLEDRK) or mutant (RLLGRSA**A**YSYLEDRK) peptide in presence of kinase buffer (40 mM Tris-HCl, 7.5 mM MgCl_2_) containing 40uM cold ATP (Sigma) and 500 μCi/μmol ATPγ^32^P (Hartmann Analytic). After removal of beads, samples were loaded on TLC Silica gels 60G F254 (Sigma) and separated using solvent (EthylAcetate, Acetic Acid, Pyridin, H2O 5:5:1:3) and dried overnight. Plates were exposed overnight to a storage phosphor screen (GE Healthcare). Screens were scanned in a Typhoon FLA 9000 system (GE Healthcare).

### Statistics and reproducibility

GraphPad Prism (v.10) was used to determine the p values and perform statistical analyses, as well as for graphical representation. Precise p values are shown in the figures. Student’s t-tests were used for comparison between two groups. If multiple groups were presented in a graph, one-way analysis of variance (ANOVA) was performed followed by the post-hoc multiple comparison. Bar graphs indicate mean ± sd. Boxplots were generated in Tukey style (1.5xIQR).

## Data availability

Data supporting the findings of this study are available within the article, its Extended Data and Supplementary table files. MS data are available on PRIDE. ImageJ custom macros are available upon request. Any additional information and wanted data are available from the corresponding author upon reasonable request.

## Extended Data

**Extended Data Fig. 1.**
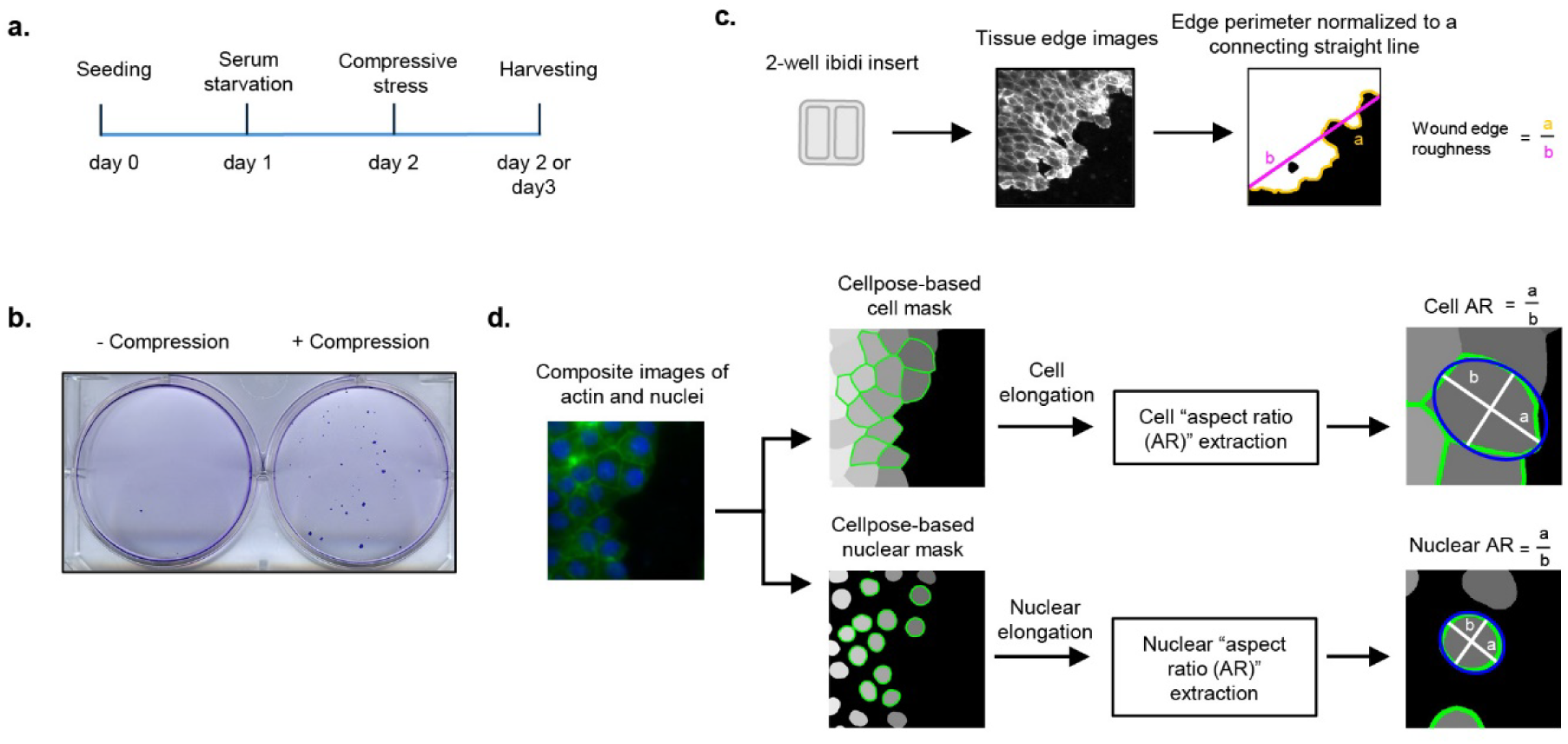
Compressive mechanical stress triggers morphological rearrangement in NMuMG cells. **a.** Experimental outline of compressive stress application on the transwell platform. **b.** NMuMG cells were exposed (+) or not (-) to compression and allowed to migrate for 24 hours through narrow 8 µm transwell pores. Transmigrated cells adhere to the bottom of the 6-well plate form colonies, which were stained with crystal violet. Representative images of colonies are shown. **c.** Wound edge roughness quantification. Images of the tissue edges obtained with ibidi chambers are converted to a mask image, and the perimeter of the edge is divided by a straight line connecting two edge points. **d.** Schematic workflow to quantify the nuclear and cell morphological changes.

**Extended Data Fig. 2.**
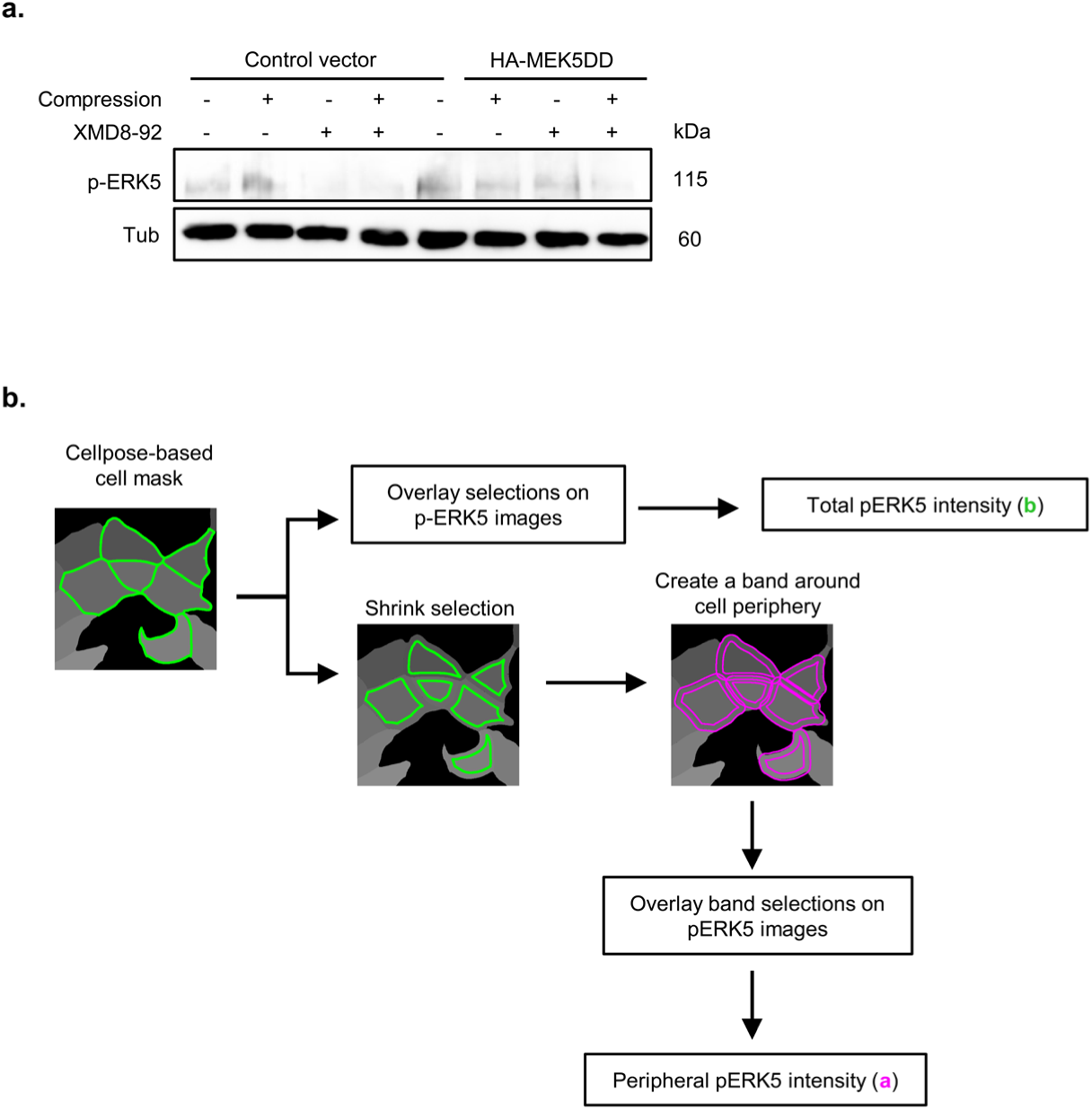
Compression leads to ERK5 activation and accumulation at cell-cell contacts. **a.** Western blot demonstrating increased p-ERK5 levels upon overexpression of HA-MEK5DD. Where indicated, the cells were compressed (+) or not (-) for 3 hours, either in the presence (+) or absence (-) of XMD8-92 **b.** Workflow illustrating quantification of total and peripheral fluorescence intensity of p-ERK5.

**Extended Data Fig. 3.**
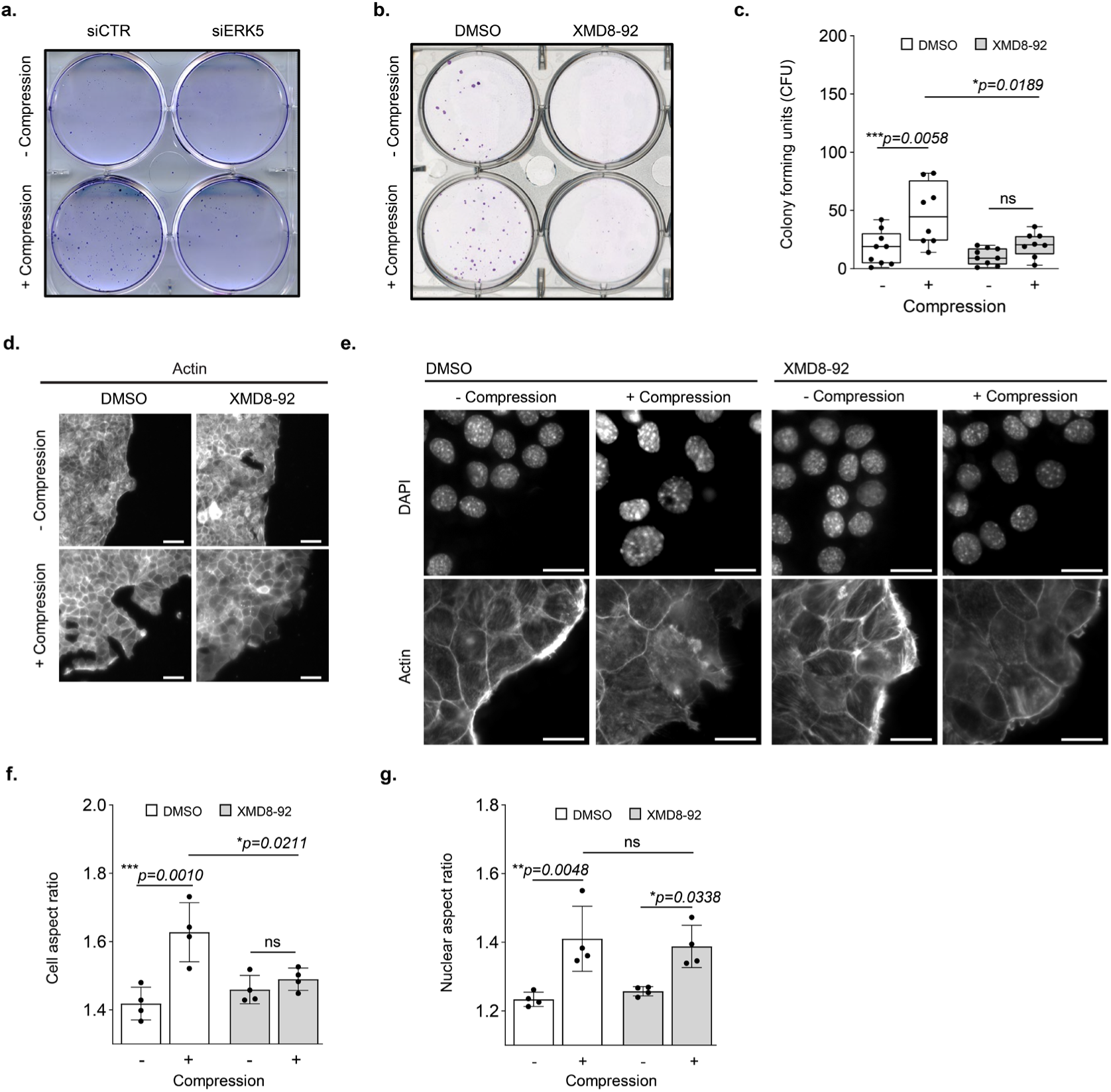
ERK5 is required for the invasive phenotype triggered by compressive pressure. **a.** and **b**. crystal violet staining of transmigrated NMuMG colonies exposed (+) or not (-) to compression and treated with siRNA controls (siCTR) or oligos targeting ERK5 (siERK5) (**a**), or upon ERK5 inhibition with XMD8-92 or the DMSO-solvent control (**b**). **c.** Quantification of colony forming units (CFU) of **b**. N=8; Statistical analysis was performed using ANOVA followed by Tukey’s multiple comparison test; ***p<0.001; p<0.05, considered significant. **d.** Invading NMuMG cells growing in ibidi chambers treated with XMD8-92 or for control with DMSO, were exposed (+) or not (-) to compression. After 24 hours, the cells were stained with phalloidin and filamentous actin visualized by microscopy. Scale bar: 50 μm. **e.** Representative images of NMuMG cells with (+) or without (-) compression were stained with DAPI to visualize the nuclei and phalloidin to probe filamentous actin. As indicated, the cells were treated with XMD8-92 or DMSO for control. Scale bar: 20 μm. **f.** Cell shape changes were quantified as the increase in the cell aspect ratio after Cellpose-based cell segmentation. N=4; ANOVA and Tukey’s multiple comparison; ***p<0.001; p<0.05, considered statistically significant. **g.** Morphological changes of nuclei are quantified as the increase in nuclear aspect ratio after Cellpose-based nuclear segmentation. N=4; ANOVA with Tukey multiple comparison; ***p<0.001; p<0.05, considered statistically significant. Note that upon compression, ERK5 activity is required for actin rearrangement and cell-cell contact dissolution.

**Extended Data Fig. 4.**
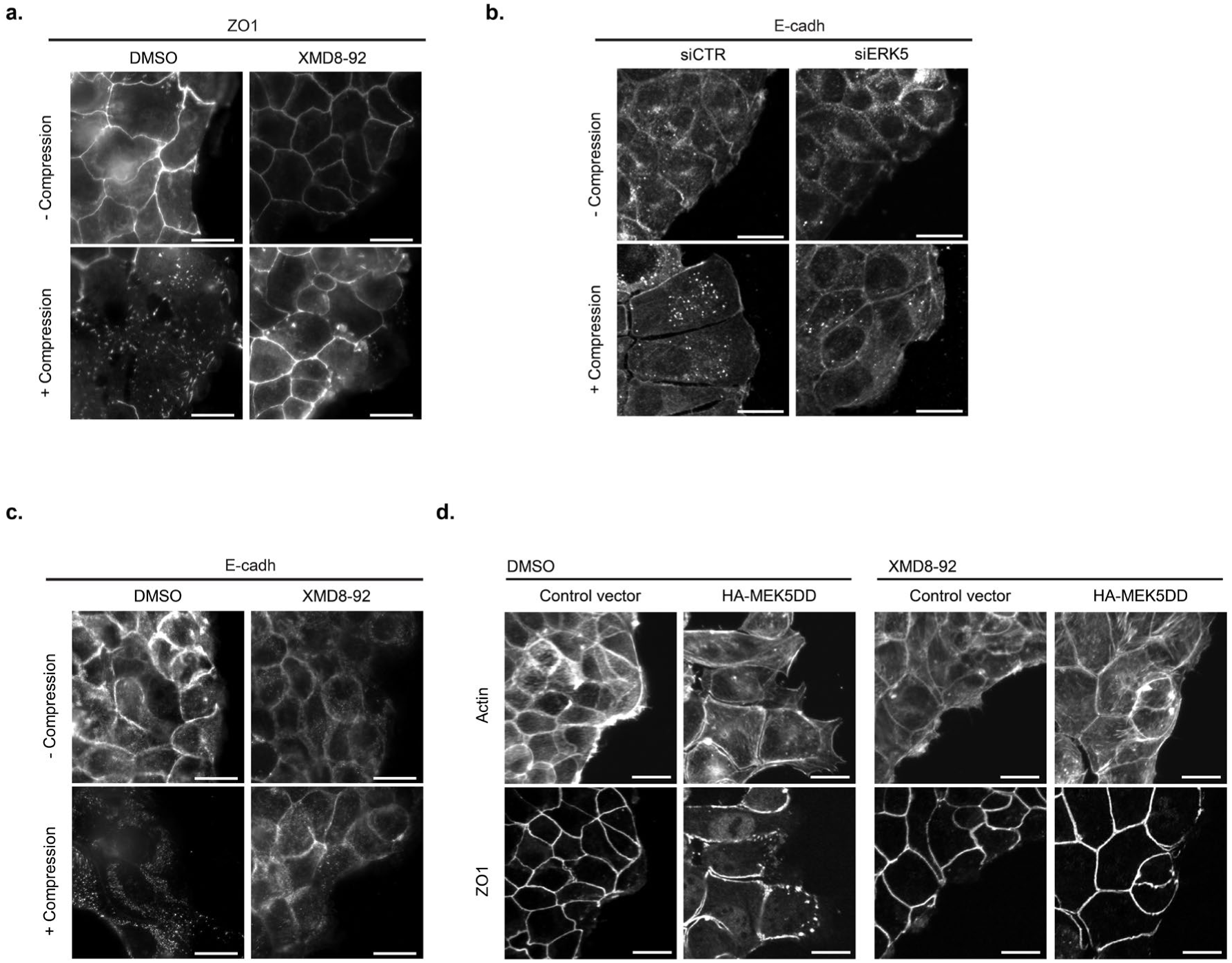
ERK5 is required for morphological changes triggered by compressive pressure. **a – c.** NMuMG cells exposed (+) or not (-) to compression were treated with XMD8-92 or for control with DMSO (**a**, **c**), or with siRNA controls (siCTR) or sioligos targeting ERK5 (siERK5) (**b**). Cells were then stained with phalloidin to visualize filamentous actin and antibodies against E-cadherin (**a**, **b**) or ZO1 (**c**). Scale bar: 20 μm. ERK5 activity is required for compression-induced dissolution of tight and adherens junctions as well as E-cadherin internalization. **d.** Confocal images of actin and ZO1 in NMuMG cells transfected with a control vector or a plasmid overexpressing HA-MEK5DD to constitutively activate ERK5. As indicated the cells were treated with the ERK5 inhibitor XMD8-92 or for control DMSO. Scale bar: 20 μm. Note that HA-MEK5DD overexpression promotes an invasive morphology even in the absence of compression.

**Extended Data Fig. 5.**
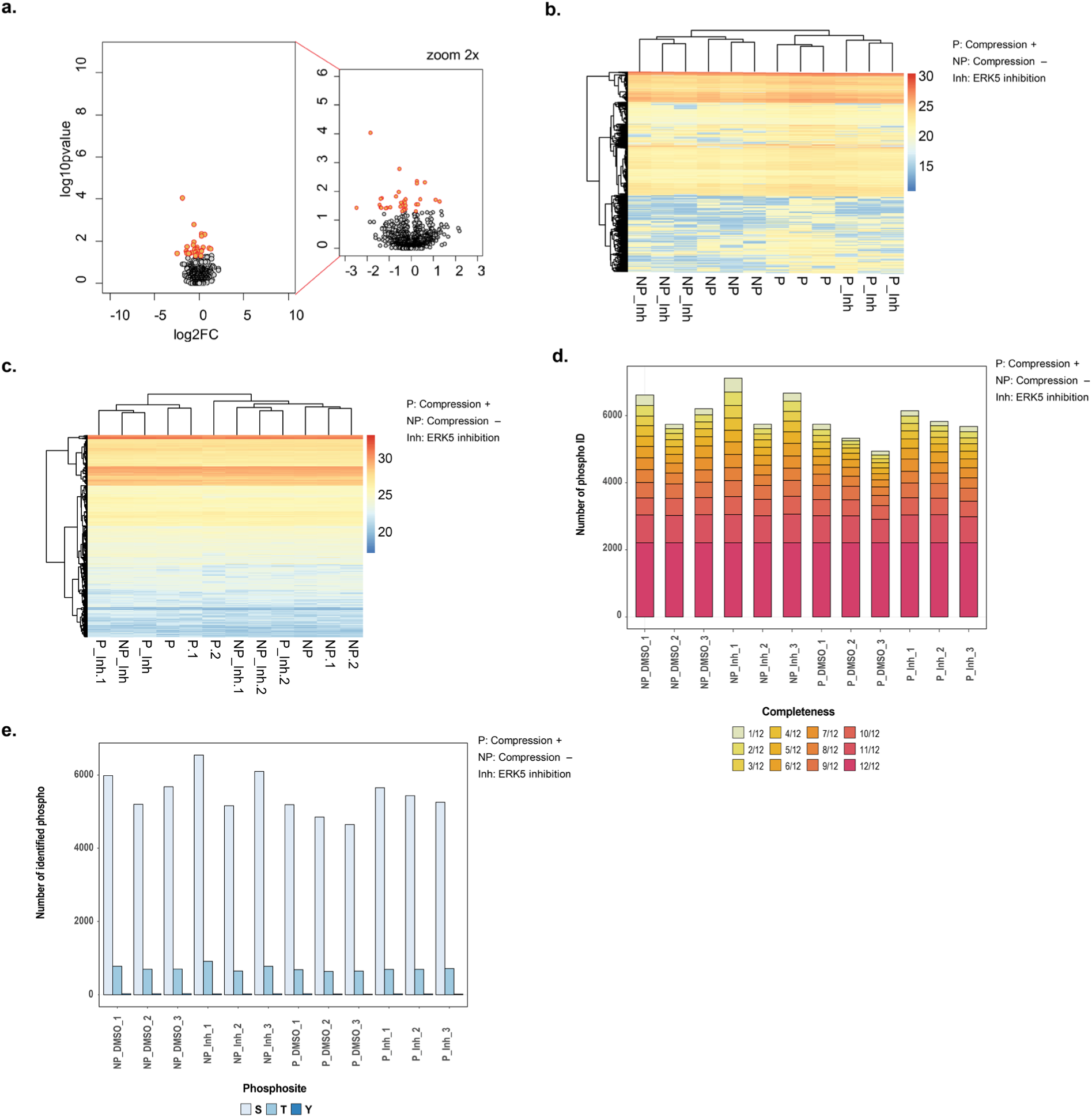
ERK5 does not regulate global protein levels upon compression. **a.** Volcano plot showing differential protein levels in compression versus no-compression comparison. Only few proteins change their level upon compression (p-adj. ≤ 0.05, log2FC>1.0), highlighting that the compressive stress response is mainly driven by phosphorylation changes. **b.** and **c**. Data clustering of the phosphoproteome (**b**) or total proteome (**c**). P: compression, NP: no compression. The addition of XMD8-92 is indicated (Inh), the number indicates biological repeats. **d.** Number of phosphopeptides identified in the phosphoproteomic experiment. **e.** Number of phosphopeptides identified in the phosphoproteomic experiment with a distribution of serine (S), threonine (T) and tyrosine (Y) phosphosites.

**Extended Data Fig. 6.**
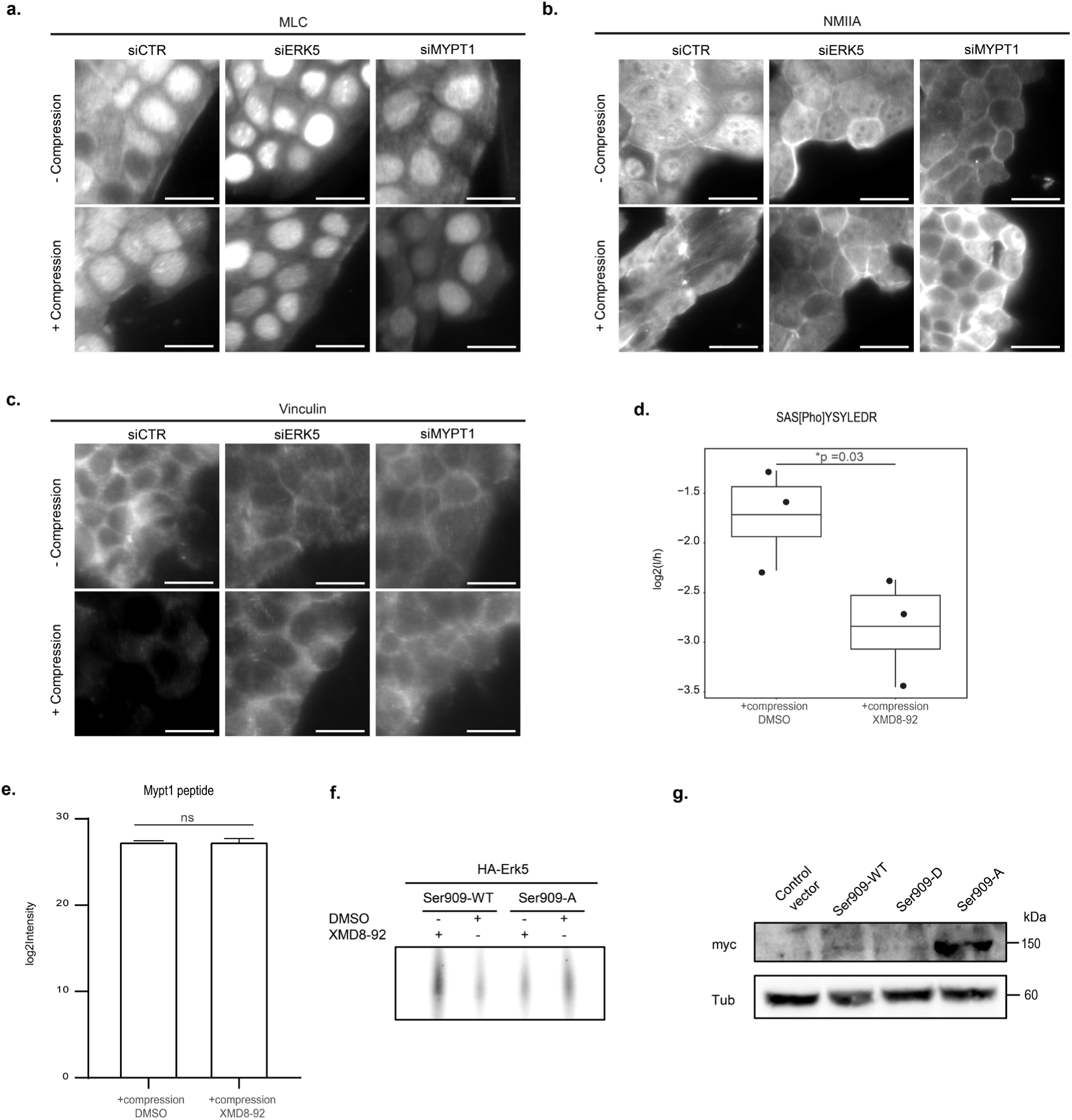
ERK5 phosphorylates MYPT1 on Ser909, thereby decreasing peripheral tension. **a**. **– c**: NMuMG cells exposed (+) or not (-) to compression were treated with siRNA controls (siCTR) or sioligos targeting ERK5 (siERK5) or MYPT1 (siMYPT1). Cells were stained for total MLC (**a**), non-muscle myosin IIA heavy chain (**b**), or the cell tension marker vinculin (**c**), and visualized by immunofluorescence. Scale bar: 20 µm. **d.** Quantification Ser909 phosphorylation of MYPT1 using PRM analysis. NMuMG cells were compressed in the presence or absence of XMD8-92 (inhibitor). N=3; One-tailed t-test; ***p<0.001; p<0.05, considered statistically significant. **e.** Analysis of total MYPT1 levels in NMuMG cells compressed in the presence or absence of XMD8-92 (inhibitor) using phosphoproteomic analysis. N=3; One-tailed t-test; ***p<0.001; p<0.05, considered statistically significant. **f.** HA-tagged ERK5 was immobilized on HA-beads and incubated in the presence of γ^32^P-ATP with a peptide encompassing the Ser909 phosphorylation site of MYPT1, either as wild-type (S909-WT) or the S909 alanine mutant (S909-A). As indicated, the *in vitro* kinase assay was performed in the presence of XMD8-92 or DMSO for control. **g.** NMuMG cells grown in ibidi chambers were reverse-transfected with an empty control vector or a vector overexpressing myc-tagged wild-type MYPT1 (Ser909-WT), or MYPT1 mutants changing S909 either to a phosphomimicking glutamic acid (Ser909-D) or a non-phosphorylatable alanine (Ser909-A) residue. Expression levels were visualized by immunoblotting with myc-antibody (upper blot). An antibody against tubulin (Tub) controls equal loading. While Ser909-A is overexpressed, Ser909-WT and Ser909-D are present at comparable levels.

**Extended Data Table 1.**

| Epitope | Company | Catalogue number |
| --- | --- | --- |
| GFP | Roche Diagnostics | 11814460 001 |
| ERK5 | Cell Signaling | #3372 |
| pERK5 (T218/Y220) | Santa Cruz | sc-135760 |
| E-Cadherin | BD Biosciences | 610181 |
| ZO1 | Cell Signaling | #2847 |
| Tubulin | Sigma Fluka Aldrich | T5168 |
| GAPDH | Sigma Fluka Aldrich | G8795 |
| MYPT1 | Cell Signaling | #2634 |
| pMYPT1 (T853) | Cell Signaling | #4563 |
| pMYPT1 (S909) | Ubiquigent | 68-0046-100 |
| MLC | Thermo Fisher | PA5-102917 |
| pMLC | Abcam | 2480 |
| Vinculin | Sigma Fluka Aldrich | V9131 |
| 9E10 (Myc) | Thermo Fisher | MA1-980 |
| NMIIA | Biolegend | 909801 |
| Goat Anti-Rabbit IgG HRP | Biorad Life Science Group | 170-6515 |
| Goat Anti-Mouse IgG HRP | Biorad Life Science Group | 170-6516 |
| Alexa-Fluor 488 Anti-Mouse | Thermo Fisher | A11029 |
| Alexa-Fluor 633 Anti-Rabbit | Thermo Fisher | A21070 |
| Alexa-Fluor 647 Anti-Rabbit | Thermo Fisher | A21245 |

## References

1. Hoffman, B., Grashoff, C. & Schwartz, M. Dynamic molecular processes mediate cellular mechanotransduction. Nature 475, 316–323 (2011).

2. Jaalouk, D., Lammerding, J. Mechanotransduction gone awry. Nat Rev Mol Cell Biol 10, 63–73 (2009).

3. Linke, J.A., Munn, L.L. & Jain, R.K. Compressive stresses in cancer: Characterization and implications for tumour progression and treatment. Nature Reviews Cancer 24, 768–791 (2024).

4. Heisenberg, Carl-Philipp et al. Forces in Tissue Morphogenesis and Patterning. Cell 153, 948 – 962 (2013).

5. Shroff, N.P. et al. Proliferation-driven mechanical compression induces signalling centre formation during Mammalian Organ Development. Nature Cell Biology 26, 519–529 (2024).

6. Tse, J. M., et al. Mechanical compression drives cancer cells toward invasive phenotype. Proc. Natl Acad. Sci. USA 109, 911–916 (2012).

7. Fernández-Sánchez, M. et al. Mechanical induction of the tumorigenic β-catenin pathway by tumour growth pressure. Nature 523, 92–95 (2015).

8. Kim, B.G. et al. Compression-induced expression of glycolysis genes in CAFs correlates with EMT and angiogenesis gene expression in breast cancer. Communications Biology 2, 313 (2019).

9. Kalli, M. et al. Solid stress-induced migration is mediated by GDF15 through Akt pathway activation in pancreatic cancer cells. Scientific Reports 9, 978 (2019).

10. Hadi T. Nia et al. Physical traits of cancer. Science 370, eaaz0868 (2020).

11. Butcher, D.T., Alliston, T. & Weaver, V.M. A tense situation: Forcing tumour progression. Nature Reviews Cancer 9, 108–122 (2009).

12. Mishra, R. et al. Protein kinase C and calcineurin cooperatively mediate cell survival under compressive mechanical stress. Proc. Natl Acad. Sci. USA 114, 13471–13476 (2017).

13. Truman, A.W. et al. Expressed in the yeast S*accharomyces cerevisiae*, human ERK5 is a client of the hsp90 chaperone that complements loss of the SLT2P (MPK1P) cell integrity stress-activated protein kinase. Eukaryotic Cell 5, 1914–1924 (2006).

14. Pereira, D.M., Rodrigues, C.M.P. Targeted avenues for cancer treatment: The MEK5–ERK5 signaling pathway. Trends in Molecular Medicine 26, 394–407 (2020).

15. Chen, Y. et al. Fluid Shear Stress Stimulates Big Mitogen-activated Protein Kinase 1 (BMK1) Activity in Endothelial Cells: Dependence on tyrosine kinases and intracellular calcium. Journal of Biological Chemistry 274, 143–150 (1998).

16. Kim, M., et al. Laminar flow activation of ERK5 protein in vascular endothelium leads to atheroprotective effect via NF-E2-related factor 2 (Nrf2) activation. Journal of Biological Chemistry 287, 40722–40731 (2012).

17. Pavan, S. et al. A kinome-wide high-content Sirna screen identifies MEK5–ERK5 signaling as critical for breast cancer cell EMT and metastasis. Oncogene 37, 4197–4213 (2018).

18. M. Angela Nieto. Epithelial Plasticity: A Common Theme in Embryonic and Cancer Cells. Science 342, 1234850 (2013).

19. Simões, A. et al. Aberrant MEK5/ERK5 signalling contributes to human colon cancer progression via NF-*κ*B activation. Cell Death & Disease 6, e1718 (2015).

20. Cronan, M. et al. Defining MAP3 kinases required for MDA-MB-231 cell tumor growth and metastasis. Oncogene 31, 3889–3900 (2012).

21. Mehta, P. et al. MEK5 overexpression is associated with metastatic prostate cancer, and stimulates proliferation, MMP-9 expression and invasion. Oncogene 22, 1381–1389 (2003)

22. Xu, Q. et al. The extracellular-regulated protein kinase 5 (ERK5) enhances metastatic burden in triple-negative breast cancer through focal adhesion protein kinase (FAK)-mediated regulation of cell adhesion. Oncogene 40, 3929–3941 (2021).

23. Gjorevski, N. & Nelson, C.M. Integrated morphodynamic signalling of the mammary gland. Nat Rev Mol Cell Biol 12, 581–593 (2011).

24. Ewald, A.J. et al. Mammary collective cell migration involves transient loss of epithelial features and individual cell migration within the epithelium. Journal of Cell Science 125, 2638–2654 (2012).

25. Nguyen, DA.D., Neville, M.C. Tight Junction Regulation in the Mammary Gland. J Mammary Gland Biol Neoplasia 3, 233–246 (1998).

26. Nia, H. et al. Solid stress and elastic energy as measures of tumour mechanopathology. Nature Biomedical Engineering 1, 0004 (2017).

27. A. J. Lomakin et al. The nucleus acts as a ruler tailoring cell responses to spatial constraints. Science 370, eaba2894 (2020).

28. Valeria Venturini et al. The nucleus measures shape changes for cellular proprioception to control dynamic cell behavior. Science 370, eaba2644 (2020).

29. Andrea K., Ferenc E. & Beáta L. Myosin phosphatase: Unexpected functions of a long-known enzyme, BBA - Molecular Cell Research 1866, 2–15 (2019).

30. Vicente-Manzanares M. et al. Non-muscle myosin II takes centre stage in cell adhesion and migration. Nat Rev Mol Cell Biol 10, 778–790 (2009).

31. Charras G. et al. Tensile Forces and Mechanotransduction at Cell–Cell Junctions. Current Biology 28, R445–R457 (2018).

32. Mayra D. A., et al. Regulation of Myosin Light-Chain Phosphatase Activity to Generate Airway Smooth Muscle Hypercontractility. Front. Physiol. 11, 701 (2020).

33. Mukta K., et al. Reconstituted Human Myosin Light Chain Phosphatase Reveals Distinct Roles of Two Inhibitory Phosphorylation Sites of the Regulatory Subunit, MYPT1. Biochemistry 53, 2701−2709 (2014).

34. Xonia C., et al. Multifunctional role of Erk5 in multiple myeloma. Blood 105, 11 (2005).

35. DuFort, C., Paszek, M. & Weaver, V. Balancing forces: architectural control of mechanotransduction. Nat Rev Mol Cell Biol 12, 308–319 (2011).

36. Guilluy, C. et al. Isolated nuclei adapt to force and reveal a mechanotransduction pathway in the nucleus. Nature Cell Biology 16, 376–381 (2014).

37. Kular J., et al. A Negative Regulatory Mechanism Involving 14-3-3ζ Limits Signaling Downstream of ROCK to Regulate Tissue Stiffness in Epidermal Homeostasis. Developmental Cell 35, 759 – 774 (2015).

38. Campàs, O., Noordstra, I. & Yap, A.S. Adherens junctions as molecular regulators of emergent tissue mechanics. Nat Rev Mol Cell Biol 25, 252–269 (2024).

39. Rooij, J. d., et al. Integrin-dependent actomyosin contraction regulates epithelial cell scattering. The Journal of Cell Biology 171, 153–164 (2005).

40. Hino N., et al. ERK-Mediated Mechanochemical Waves Direct Collective Cell Polarization. Developmental Cell 53, 646–660.e8 (2020).

41. Bastianello G., et al. Cell stretching activates an ATM mechano-transduction pathway that remodels cytoskeleton and chromatin. Cell Reports 42, 113555 (2023).

42. Elosegui-Artola A., et al. Force Triggers YAP Nuclear Entry by Regulating Transport across Nuclear Pores. Cell 171, 1397 - 1410.e14 (2017).

## References

43. Edelstein, A., et al. Computer control of microscopes using μManager. Curr. Protoc. Mol. Biol. 92, 14.20.1–14.20.17 (2010).

44. Schneider, C. A., Rasband, W. S. & Eliceiri, K. W. NIH Image to ImageJ: 25 years of image analysis. Nature Methods 9, 671–675 (2012).

45. Stringer, C., et al. Cellpose: a generalist algorithm for cellular segmentation. Nature Methods 18, 100–106 (2021).

46. Cox, J., Mann, M. MaxQuant enables high peptide identification rates, individualized p.p.b.-range mass accuracies and proteome-wide protein quantification. Nature Biotechnology 26, 1367–1372 (2008).

47. MacLean B., et al. Skyline: an open source document editor for creating and analyzing targeted proteomics experiments. Bioinformatics 26, 966–968 (2010).

